# Edition of complex gene families in tobacco with GoldenBraid 4.0, a multipurpose web-based platform for plant genome engineering

**DOI:** 10.1101/2020.10.06.327841

**Authors:** Marta Vazquez-Vilar, Víctor Garcia-Carpintero, Sara Selma, Joan M Bernabé-Orts, Javier Sanchez-Vicente, Blanca Salazar-Sarasua, Arianna Ressa, Carmine de Paola, María Ajenjo, Asun Fernández-del-Carmen, Antonio Granell, Diego Orzáez

## Abstract

CRISPR/Cas ability to target several loci simultaneously (multiplexing) is a game-changer in plant breeding. Multiplexing not only accelerates trait pyramiding but also can unveil traits hidden by functional redundancy in polyploid crops. Furthermore, multiplexing enhances dCas-based programmable gene expression and enables cascade-like gene regulation. However, multiplex constructs comprising tandemly arrayed gRNAs are difficult to assemble, this hampering more widespread use. Here we present a comprehensive upgrade of the popular cloning platform GoldenBraid (GB), in which, on top of its classical multigene cloning software, we integrate new assembly tools for two-dimensions gRNA multiplexing with both Cas9 and Cas12a, using the gRNA-tRNA-spaced and the gRNA unspaced approaches, respectively. As functional validation, we show, among others, the assembly of up to 17 tandemly-arrayed gRNAs constructs against a subset of the Squamosa-Promoter Binding Protein-Like (SPL) gene family in tobacco. With these constructs we generated a collection of Cas9-free SPL mutants harboring up to 9 biallelic mutations in a single generation. The functionality of GB-assembled dCas9 and dCas12a-based CRISPR activators and repressors using single and multiplexing gRNAs is also validated. With the incorporation of the new CRISPR tools and part’s collection, GB4.0 turns an unprecedentedly comprehensive open platform for plant genetic engineering.

## INTRODUCTION

Genome editing tools based on CRISPR/Cas enzymes have rapidly become central players in modern plant breeding due to their unprecedented precision and the simplicity of its molecular machinery, which involves only a constant endonuclease element and an easy-to-program RNA guide. This simplicity explains its widespread use in many labs and small companies which have no access to previous editing tools. For most species including many crops, agrobacterium-mediated transformation is the method of choice for the delivery of editing machinery into the plant. In this case, a transgenic intermediary plant is produced as a consequence of the stable integration of the CRISPR/Cas-containing transgene in the genome, which can be later segregated to create transgene-free edited plants. Although the employment of intermediary transgenic steps could eventually pose regulatory hurdles, the use of transgene constructs also brings along all the advantages of multigene engineering, which can expand enormously the capabilities of CRISPR/Cas not only as site-specific nuclease for mutagenesis but also as a programmable tool for additional genome engineering applications as base editing (Shimatani et al., 2017; Zong et al., 2017; Kang et al., 2018), epigenome editing (Papikian et al., 2019), etc. In addition, multigene constructs ensure a physical linkage between the different gene elements employed in the editing process (i.e. selection marker, nuclease and gRNAs), facilitating segregation of transgenes and consequently the generation of transgene-free plants. Furthermore, it is unclear that the final shape of the regulatory framework in many countries will make distinctions among the different delivery procedures provided that the final plant or plant products is devoid of exogenous DNA sequences.

Recently, synthetic biology-inspired modular cloning strategies based on Type IIS restriction enzymes (Engler et al., 2009) have expanded our capacities for building combinatorial and multigene DNA constructs in binary vectors. Several modular cloning systems have been described (Sarrion-Perdigones et al., 2011; Weber et al., 2011), many of which follow the standard syntax earlier proposed for this type of systems (Patron et al., 2015). CRISPR construct-making benefits from the capacity of Golden Gate cloning to assemble large constructs containing several repeated elements. This feature facilitates the cloning of tandem gRNA constructs targeting several loci simultaneously, a strategy known as multiplexing. The importance of CRISPR multiplex editing in plant breeding cannot be underscored, as it paves the way for pyramiding favourable independent traits at unprecedented speed (Zhang et al., 2020). This has been recently exemplified with the fast domestication of wild tomato (Zsögön et al., 2018), or the fast adaptation to urban agriculture (Kwon et al., 2020). In addition, multiplexing has the ability to uncover valuable traits which have remained elusive to breeding due to redundancy in large gene families. This is more evident in polyploid plants, which account for some of the most important crop species. Remarkable examples are low gluten wheat (Sánchez-León et al., 2018), glyco-engineered *N. benthamiana* plants (Jansing et al., 2019) or semi-dwarf rapeseed with increased yield (Zheng et al., 2020).

Our lab has established over the years a renowned cloning system for plant bioengineering named GoldenBraid (GB) (Sarrion-Perdigones et al., 2011; Sarrion-Perdigones et al., 2013). The most distinctive features of GB are the SynBio-inspired standardization of its DNA parts and the exchangeability of all GB-made constructs. Thus, owing to its iterative cloning strategy, any pair of GB constructs can be straightforwardly assembled together with a golden gate reaction (Engler et al., 2009), greatly simplifying the creation of complex multigene constructs. Successive versions of the system have extended its usability and its applications. Notably, GB version 3.0 incorporated a dedicated web that serves both as a software-assisted cloning tool as well as a repository of plant genetic elements comprising > 600 public standard DNA parts including promoter regions, CDS, terminators, but also exchangeable transcriptional units and multigene constructs for e.g. conditional transgene expression, selection markers, etc. (Vazquez-Vilar et al., 2017). Each GB element is documented by a standard datasheet, which often incorporates functional (experimental) characterization.

Here we take advantage of the well-established GB platform to create a new multipurpose bioengineering resource that exploits the synergies between CRISPR tools on one side, and modular cloning and Synthetic Biology on the other side to facilitate genome engineering to the plant biotech community. The new GB4.0 (genome edition) platform contains an expanded database and a public repository that incorporates (i) all elements required for Cas9 and Cas12a single-guide and multiplex editing, plus (ii) new DNA parts and constructs required for dead Cas9-based programmable gene regulations, including also DNA elements for multiplex targeting. Furthermore, we have enriched our GB software tools so that all new cloning procedures, including all new multiplexing tools are fully software-assisted, generating output files with detailed laboratory protocols, annotated GenBank constructs, and vSBOL-informed datasheets. As a general quality control, we have been experimentally validating all the genetic elements contained in the public GB4.0 repository, and the validation results are presented here or elsewhere (Bernabé-Orts et al., 2019; Selma et al., 2019). Furthermore, taking advantage of the versatility of the cloning system, the operative limits of some of the most useful GB4.0 tools were also investigated. In a first example, we tested two large 11 and 17 gRNAs multiplex constructs used to knock-out candidate genes belonging to the MPO and SPL tobacco gene families, respectively. These experiments generated highly diverse population of T0 mutant plants, some of which accumulated up to five bi-allelic and four heterozygous mutations in as many homologue genes and leading to transgene-free T1 plants with up to 9 biallelic mutations. In a separate example, we tested the ability of dead Cas (dCas)-based programmable transcriptional activator to super-activate a strong plant promoter, yielding expression levels well above those of the constitutive 35S promoter. Noteworthy, and owing to its initial design as a community collaborative tool, GB4.0 offers all users the possibility to easily exchange their newly developed constructs/tools with the community, promoting a strongly needed cooperation and democratization of the new breeding techniques.

## RESULTS

### Software-assisted cloning of guide RNA expression-cassettes

GB cloning operates at three hierarchically organized assembly levels: Level 0 parts are basic elements such as promoters, CDSs, terminators, etc, which can be assembled using multipartite Golden Gate reactions to create level 1 elements, usually transcriptional units (TUs). Level 1 elements are combined binarily to create higher order multigene structures (named level >1 elements), following an iterative cloning pipeline. In GB4.0, we adopted the same GB assembly pipeline for the design of multigene constructs required for genome engineering, and new dedicated webtools were added to fulfil the specific requirements of CRISPR constructs. The new tools were conceived to provide a centralized user-friendly gateway for the design of CRISPR/Cas constructs, both for genome editing and for other expanded applications as gene regulation. All GB-software tools provide Genbank output files next to a vSBOL-based representation of the generated construct and a detailed protocol.

The basic GoldenBraid CRISPR/Cas genome editing construct comprises at least three level 1 transcriptional units (TUs): a TU containing the guide RNA(s), a Cas9-expressing TU, and a third unit usually encoding a selection marker TU. From a modular cloning perspective, the most demanding element is the gRNA unit. For the expression of gRNAs, the RNA pol III promoters, which specifically transcribe small nuclear RNAs in the cell, are preferably used. The gRNA TU can be designed to express just one gRNA (single RNA strategy), or engineered to contain several tandemly arrayed gRNAs under the control of a single (usually pol III) promoter (polycistronic strategy). In both cases, subsequent GB cloning iterations allow the assembly of additional gRNA TUs (single or polycistronic) to expand the multiplexing capacity. Since gRNA assembly is usually the most cumbersome step in CRISPR construct building, specific software-assisted GB cloning tools were developed for the assembly of single or polycistronic gRNA arrays, both for Cas9 and Cas12a (see Fig. 1 and https://gbcloning.upv.es/tools/grna/). Prior to gRNA assembly, users need to find convenient protospacers for targeting the desired gene(s), using for this purpose external tools as those suggested as links in the GB webpage (i.e. Benchling (https://www.benchling.com/), CRISPOR (Concordet and Haeussler, 2018), CINDEL (Kim et al., 2017), CRISPR-DT (Zhu and Liang, 2019)).

**Figure 1.**
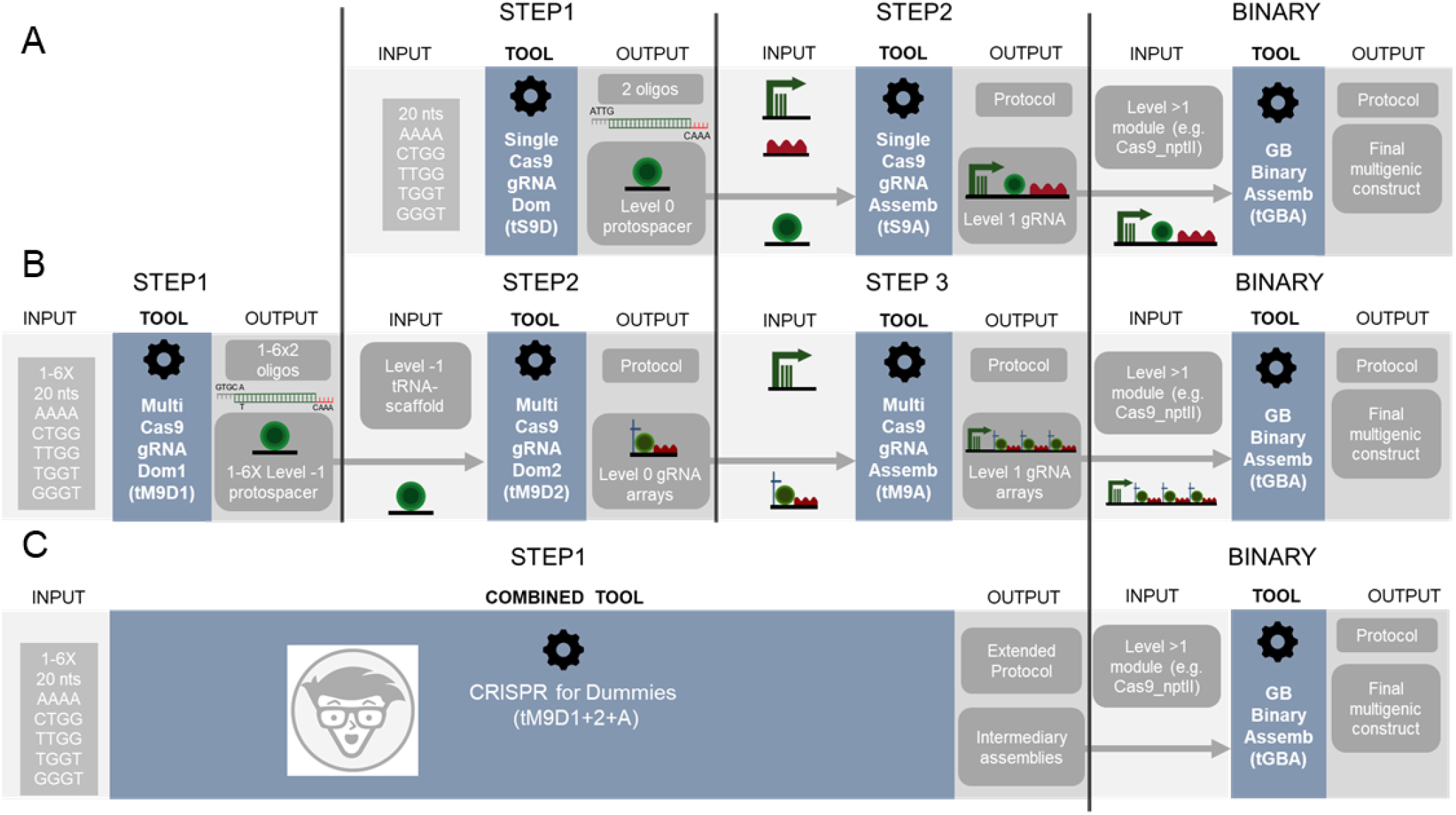
Flowchart of the software-assisted cloning strategies of Cas9 single and multiplexing guide RNA expression cassettes with GoldenBraid. A) Cas9 single gRNAs are assembled in two steps via the *Single Cas9 gRNA Domesticator* (*tS9D*) that takes as input a 20 nts protospacer sequence and the *Single Cas9 gRNA Assembler* (*tS9A*) that creates a Level 1 construct with a PolIII promoter, hybridized primers including the protospacer sequence and the corresponding overhangs, and the Cas9 scaffold. B) Cas9 multiplexing gRNAs are assembled with the use of three consecutive tools: the *Multi Cas9 gRNA Domesticator 1* (*tM9D1*) that designs partially complementary oligonucleotides that include the protospacer sequence, the *Multi Cas9 gRNA Domesticator 2* (*tM9D2*) that creates Level 0 tRNA-protospacer-scaffold units using the hybridized primers and the corresponding Level -1 vectors and the *Multi Cas9 gRNA Assembler* (*tM9A*) that creates Level 1 gRNA arrays. C) The CRISPR for Dummies tool takes 1 to 6 protospacers as input and generates Level 1 gRNA arrays in a single step.

For the assembly of single gRNA constructs for Cas9 editing, a two-steps procedure was established (Fig. S1). In a first step, the user-selected protospacer sequence (20 nucleotides) is introduced as input into the *Single Cas9_gRNA domesticator tool* (*tS9D*). The domesticator produces an output consisting in a pair of overlapping oligonucleotides containing the protospacer sequence itself, plus the appropriate cloning overhangs. It is important to mention that the custom oligos designed by the CRISPR domesticator tool are the only elements external to the GB collection that will be required for any Cas9 CRISPR construct to be made. All the remaining elements are provided. A second step, assisted by the *Single Cas9_gRNA Assembler tool* (*tS9A*) takes the pair of oligonucleotides as input and combines them in a BsaI multipartite reaction (step 2) with a Pol III promoter and the gRNA scaffold, both elements provided in the collection (Fig. 1A). This second webtool provides also a detailed protocol of the assembly reaction.

GB also includes Cas12a in its current toolbox. The Cas12a nucleases group has special assembly requirements derived from its special features: first, opposite to Cas9, the native constant elements of the RNA scaffold of Cas12a are located at the 5’end of the structure, whereas the protospacer is located at the 3’end. Second, protospacers length range between 20 and 23 nucleotides, and third, Cas12a is able to self-process a polycistronic RNA, a feature that Cas9 lacks. Consequently, independent strategies were designed for the GB cloning of Cas12 single and polycistronic gRNA constructs.

The single gRNA Cas12a design comprises two steps (Fig. S1). In the first one, a new level 0 element is designed using a dedicated design tool, the *Single Cas12a gRNA domesticator (tS12D)*. This tool takes a 20-23 nucleotides protospacer input and designs a pair of overlapping oligonucleotides containing this sequence and the appropriate overhangs for cloning. In a second step, the *Single Cas12a gRNA assembler* (*tS12A*) combines the *S12D* output with the remaining level 0 elements, namely an upstream constant element comprising the pol III promoter and the gRNA directed repeats (DRs), and the HDV ribozyme that will trim the 3’ end to expose the final nucleotide of the protospacer.

We used these tools to assemble a set of individual gRNAs both for Cas9 and Cas12a targeting both transgenic (i.e. the 35s, Nos and Mtb promoters) and endogenous sequences (i.e. XT, FT, CBP and ALS). In order to determine their editing efficiencies and to evaluate the predictive power of the ‘Rule Set 2 scoring’ (Doench et al., 2016) and CINDEL (Kim et al., 2017) for Cas9 gRNAs and Cas12a crRNAs respectively, we carried out a simple transient expression experiment in *N. benthamiana* plants. We observed a correlation coefficient of 0,82 between experimentally determined editing efficiencies and CINDEL predicted scores for Cas12a crRNAs while for Cas9 gRNAs we could not find correlation between the predicted scores and experimental efficiencies (Fig. S2).

To create polycistronic gRNA arrays for Cas9 editing, the GB pipleline uses tRNAs as spacers, which are later processed in planta by endogenous RNAses (Fig. S1). A number of designated software tools guide user through a tree-step cloning process: first, individual protospacers are domesticated and added to the database (assisted by the *Multiple Cas9_gRNA domesticator tool 1, tM9D1*); second, level 0 parts (individual gRNAs) are constructed, each comprising a tRNA, the previously domesticated protospacers and a scaffold element, all three assembled in a BsmBI reaction (*Multiple Cas9_gRNA domesticator tool 2, tM9D2*); third, a polycistronic gRNA TU is assembled combining up to six level 0 gRNAs from step 2, plus a pol III promoter (*Multiple Cas9_gRNA assembler, tM9A*). Again, all elements except the protospacer oligos are pre-made and available in the collection. Users only need to decide beforehand (step 2) the number of gRNAs (from one to six) that will conform the array, and select the level 0 parts to be created accordingly (Fig. 1B).

Multiple Cas12a gRNA design also comprises two steps (Fig. S2), assisted by the *Multiple Cas12a_gRNA domesticator (tM12D) and the Multiple Cas12a_gRNA assembler (tM12A)*. First, the *Multiple Cas12a_gRNA domesticator* takes as input two to four 20-23 nucleotides protospacer sequences and designs a tandem of scaffold-protospacer units flanked by BsmBI sites as an output. It should be noted that the output of the *tM12D* tool is a >200-500bp DNA fragment, which needs to be produced via chemical synthesis and assembled as a Level0 part. Next, the *Multiple Cas12a_gRNA assembler* combines the *tM12D* output with the remaining Level 0 elements, which are the same as those used by the *tS12A*. In this sense, Cas12 polycistronic gRNA requirements are more demanding in terms of DNA synthesis that Cas9, where only a pair of overlapping oligos was required.

Once a single or multiplexed level 1 Cas9 or Cas12a gRNA TU is created, this can be combined with other TUs using the regular GB Binary assembly tool. For convenience, the public GB collection contains a number of frequently used pre-made TUs and multigene modules, such as SpCas9_TUs, LbCas12a_TU, negative and positive selection markers, and combinations of those (*e*.*g*. GB0639, GB2234, GB1441, GB2085, etc.). Several gRNA TUs (single or polycistronic) can be combined in the same binary fashion to create large “two dimensions” multiplexing arrays. Detailed lists of recommended GB TUs and modules for each application, namely gene editing, gene activation or gene repression either with Cas9 or with Cas12a can be accessed from https://gbcloning.upv.es/tools/crispr/.

An extremely simplified version of the above mentioned Cas9 multiplexing tool was also created separately, nick-named “*Editing for Dummies*” tool (Fig. 1C, https://gbcloning.upv.es/do/crispr/cas9_multiplexing/crispr_for_dummies/). In this tool, input choices are reduced to the number (N) of arrayed gRNAs to be assembled (from one to six) and the sequences of the 20 nucleotide protospacers to be used. As an output, user’s obtain three sets of information: (i) the sequence of the 2xN oligos for synthesis; (ii) a detailed N days laboratory protocol where all GB elements to be used in the assembly are include and (iii) genbank and sbol files containing construct for general use which comprises: a multiplexed gRNA TU, a constitutively expressed hCas9, an nptII gene for negative selection, a DsRed gene for positive selection.

### GB-made multiplex constructs facilitate fast pyramiding of gene families in tobacco

As a proof of functionality, here we describe the use of GB-built Cas9 multiple gRNA constructs assayed for fast mutant pyramiding in two allotetraploid tobacco gene families, (i) the N-methylputrescine oxidase (MPO) gene family, with 6 gene members, and (ii) a subset of 14 members of the SQUAMOSA promoter binding protein-like transcription factors family. Using the GB assembly tools described above, we assembled two editing constructs, GB2484 and GB2714. Each construct comprised an nptII plant selection marker, a Cas9 TU, fluorescent DsRed as positive selection marker, and series of multiple gRNA arrays targeting MPO (Fig. 2A) and SPL (Fig. 3A) genes respectively. GB2484 construct comprised a total of 11 gRNAs distributed in two TUs of 6X and 5X gRNAs respectively, designed to target each MPO family member in three different positions (Fig. S3). On the other hand, GB2714 construct was distributed in three arrays of 6X, 5X and 6X gRNA arrays, directed to one to three different positions per target gene in the SPL subfamily (Fig. S3).

**Figure 2.**
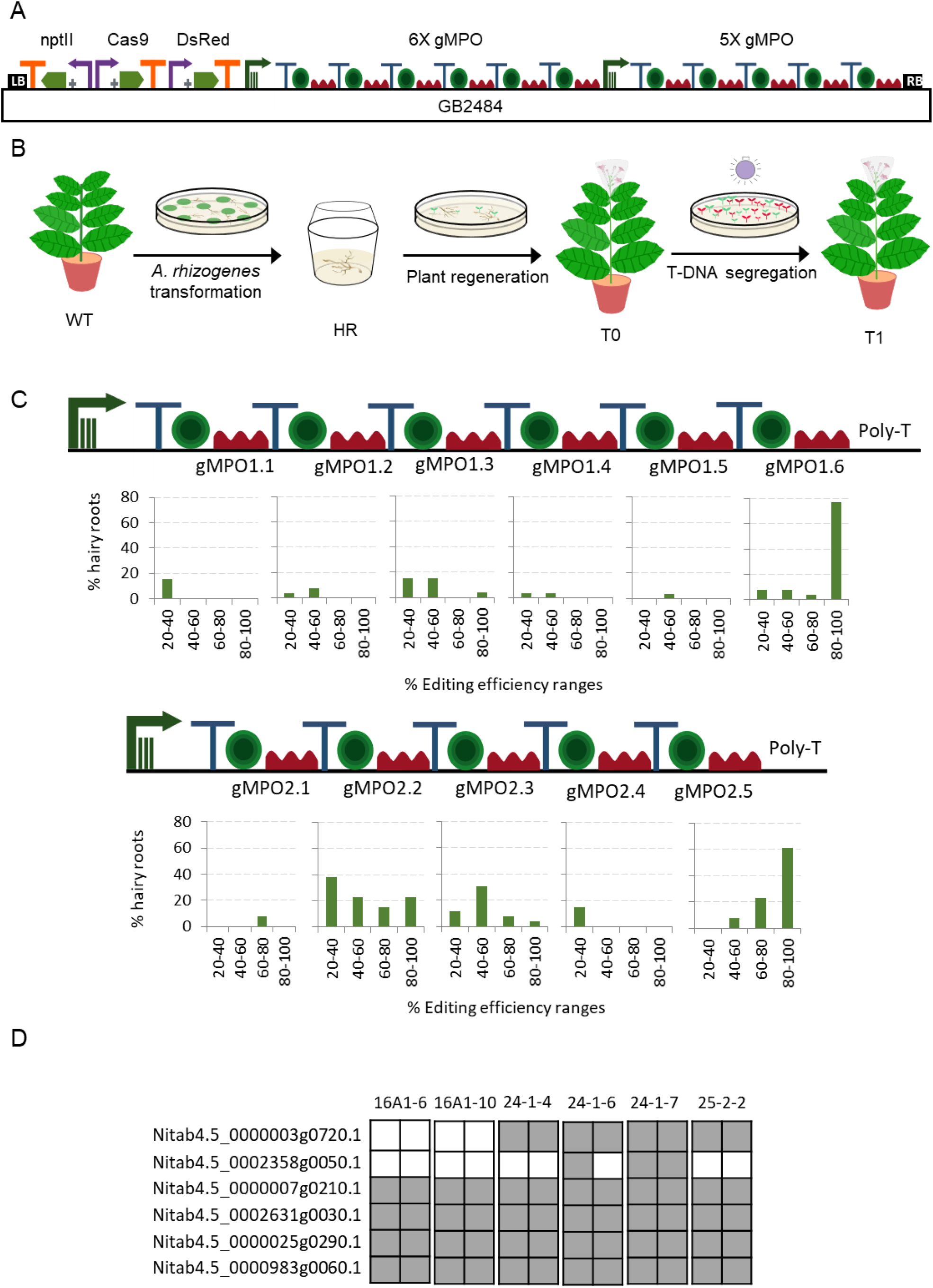
Cas9 guide RNAs editing efficiencies in tobacco hairy roots. A) Construct including 11 gRNAs (6X+5X) targeting MPO1 genes that was used for *A. rhizogenes* transformation. B) Schematic representation of tobacco transformation with *A. rhizogenes*. C) gRNA efficiency for each position in the polycistron evaluated for 13 tobacco hairy roots by jointly analyzing the data obtained for all genes targeted by the same gRNA. D) Schematic representation of a selection of TDNA-free T1 MPO edited lines. Each square represents an allele. Grey indicates edited alleles while white represents wild type alleles.

**Figure 3.**
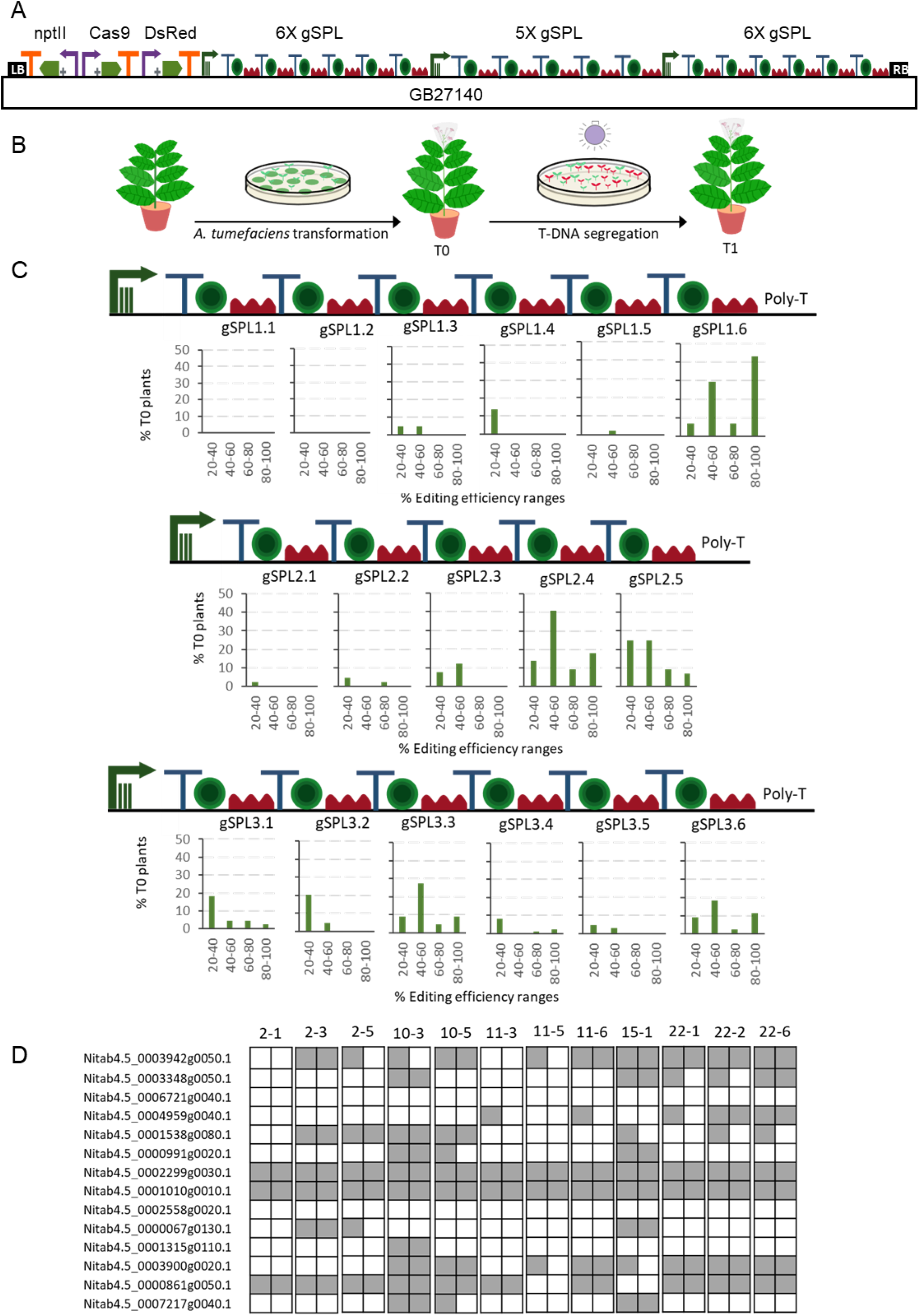
Cas9 guide RNAs editing efficiencies in tobacco stable transformation. A) Construct including 17 gRNAs (6X+5X+6X) targeting SPL genes that was used for *A. tumefaciens* transformation. B) Schematic representation of tobacco transformation with *A. tumefaciens*. C) gRNA efficiency for each position in the polycistron evaluated for 22 T0 tobacco plants by jointly analyzing the data obtained for all genes targeted by the same gRNA. D) Schematic representation of a selection of TDNA-free T1 SPL edited lines. Each square represents an allele. Grey indicates edited alleles while white represents wild type alleles.

The GB2484 MPO construct was transformed in wild type *N. tabacum* cv. K326, using *A. rhizogenes* yielding 13 hairy root explants, which were subsequently genotyped (Fig. 2B). The editing efficiencies obtained by each gRNA in each target gene are depicted in Fig. 2C. In a second transformation experiment, the GB2714 SPL construct was transformed in a tobacco plant line (157-5) of the same *N. tabacum* cv. K326 variety, which had been previously mutagenized in a different subset of genes in the SPL family (Fig. 3B). In this later case, *Agrobacterium tumefaciens* was used for transformation, and the resulting 22 T0 regenerated plants were analysed by ICE (Synthego) to estimate the editing efficiency of each individual gRNA as summarized in Fig 3C.

To better visualize possible influences of positional effects in the array, editing efficiencies estimates were grouped in four ranges, with only rates above 20% considered as positive. Mutation estimations between 40%-60% were presumed as heterozygous mutations; rates estimated above 80% were regarded as biallelic, and the remaining intermediate levels were considered as chimeric affecting one (60-80%) or both (20-40%) alleles. As expected, edition estimates in both experiments show highly variable values, with two gRNAs (gSPL1.1 and gSPL1.2) showing values below the threshold, whereas others as gMPO1.6 showed almost 80% putative biallelic rates. It is important to note that all positions in both 5X and 6X arrays yielded estimates above the 20% threshold for at least one of the gRNAs assayed, indicating that all array positions are active. However, data suggest some positional bias in editing efficiency. As can be observed, in all five polycistronic arrays assayed, biallelic mutations were recovered for targets whose gRNA was in the last position (from 5% in gSPL2.5 to 80% in gMPO1.6). In contrast, only one gRNA set in position 1 (gSPL3.1) was able to produce biallelic plants, and only in a small proportion (5%). In general, the last position turned out to be the most effective in three of the five polycistronic arrays, whereas in the remaining two arrays positions 3 and 4 were first in the ranking of efficiency.

Both MPO and SPL multiplexing strategies successfully pyramided multiple-knock outs in T0 and T1 generations, the latter easily made transgene free by DsRed (-) selection of T1 seeds grown in vitro (later tested by PCR, data not shown). The best MPO hairy roots accumulated up to 2 full (biallelic), 1 heterozygous KO genes and chimerism on the other 3 genes, whereas in T1 generation, one plant out of 6 DsRed (-) analysed showed biallelic mutations in all six MPO genes (Fig. 2D, Table S4). Unfortunately, this line still contained the Ri DNA, which was removed in T2 generation. A second T1 plant was recovered showing not R-not T-DNA having five biallelic mutations. In the case of the SPL genes, the highest mutation rates obtained in T0 corresponded to a plant having 6 biallelic and 3 heterozygous mutations. The best T-DNA-free T1 line showed up to 9 biallelic and 1 heterozygous mutations in as many SPL genes (plant SPL-10-3). As mentioned above, the present experiment was performed on top of a previously mutagenized T1 plant harbouring 6 biallelic mutations in a different subset of SPL genes, therefore the resulting genotype of SPL-10-3 comprised 15 KOs out of the 20 SPL gene family in *N. tabacum* (Fig. 3D, Table S4). While SPL T0 plants were selected based on the diversity on their mutations, MPO T0 plants were selected based on the number of edited genes. The differences between the mutations of the T1 progeny displayed on Fig. 2D and Fig. 3D reflect these differential selection processes.

### Beyond DNA cuts: GB-made dCas9 and dCas12a-based programmable regulators for gene activation and repression

GB4.0 also incorporates webtools and DNA elements for the design of Cas9 and Cas12a-based programmable transcriptional regulators (PTRs). Programmable regulators can be created by adding transcriptional regulatory domains (Repressor Domains, RDs, or Activator Domains, ADs, jointly regarded as regulatory domains, RgDs) to a nuclease-inactivated (dead) dCas, either directly as protein fusions or indirectly via a multiepitope peptide. In the latter case, multiple RgDs can be attached to the multiepitope peptide via a single chain antibody (scFv) intermediary (SunTag strategy, (Tanenbaum et al., 2014)). In both direct and indirect examples, the design of a gRNA TU remains the same as in the previously described editing constructs. The only difference in the design pipeline takes place during binary assemblies, where transcriptional units conveniently equipped with dCas-RgD or dCas-SunTag plus scFv-RgD will be incorporated instead of the regular Cas9 endonuclease. Several dCas-RgD and dCas9-SunTag fusions are available in the GB public repository, together with different scFv-RgD fusions to complement the SunTag strategy (https://gbcloning.upv.es/tools/crispr/regulation/TUs/)

A more elaborated regulatory strategy consists in introducing modified gRNAs that incorporate RNA aptamers attached to different positions of the gRNA scaffold (SAM and scRNA strategies, (Konermann et al., 2015)). We previously demonstrated that a modified scRNA strategy (dCas9-EV2.1) resulted in a potent programmable activator tool in plants, able to selectively upregulate up to 10000 folds the stress-inducible DFR promoter (Selma et al., 2019). Therefore, we developed specific GB parts and software-tools for assisting in the assembly of gRNAs for dCas9-EV2.1 strategy. Briefly, single and multiplexed (up to 3X) modified gRNA arrays for transcriptional regulation can be built stepwise using the *Multiple Cas9_gRNA Domesticator tool 1 (tM9D1*) and *Multiple regulatory Cas9_gRNA Domesticator tool 2 (tMr9D2*) tools. These tools are equivalent to the above described *tM9D1* and *tM9D2* except that now the input gRNA scaffold contains an RNA aptamer that binds MS2.

Here we tested the functionality of GB4.0-built transcriptional Cas9-based activators by over-activating the SlMtb promoter. Given the ability of CRISPR-based activators to activate DFR and other inducible promoters, we wanted to investigate here if they could also serve to increase the transcriptional levels conferred by a strong constitutive promoter. The 2 kb upstream promoter region of the *S. lycopersicum* Mtb (Metallothionein-like protein type 2B, Solyc09g010800) gene, (catalogued as GB0080) has a strong constitutive activity, reaching approximately four times that of the pNos promoter activity used as reference in GB standard measurements. Promoter Relative Transcription Activity (RTA) is estimated in *N. benthamiana* leaves using the Luciferase/Renilla dual reporter system, and normalizing the luminescence levels conferred by the test promoter with those produced by the pNos promoter in the same experimental conditions. The resulting normalized value is expressed as relative promoter units (rpu). The RTA for GB0080 promoter was earlier estimated as 4+/-1 rpu, about 1/3 of a 35S promoter (GB0030, RTA= 10 +/-2 rpu). The transcriptional over-activation of the pSlMtb was first analysed using the dCas9-EV2.1 complex combined with gRNAs at positions +41, +5, -50 and -402 (relative to the Transcription Start Site, TSS), as represented in Fig. 4A. The gRNAs were tested individually or combined in a single T-DNA as depicted. All gRNAs tested in a window of -400 to +5 relative to the TSS conferred strong activation to the promoter, with gRNA overlapping the TSS reaching maximum levels (Fig. 4C). The combination of all four gRNAs in a single T-DNA, conferred activation levels only slightly higher than those obtained by gRNA +5 acting individually. Most notably, absolute RTA levels obtained with 4X gRNAs reached record RTA levels (72 +/-8 rpus), corresponding to a 17-fold activation from pSlMtb basal levels and 7 times above CaMV35S levels used in this experiment as upper limit reference. The high activation conferred by the dCas-EV2.1 complex was also confirmed in a separate activation experiment using the same 4X gRNAs but combined with other activation domains in the GB collection (dCas9:VPR-MS2:VPR and dCas9:TV-MS2:VPR). As shown in Fig. 4D, the results confirmed earlier observations that the dCas-EV2.1 activator complex achieved the highest promoter activation rates.

**Figure 4.**
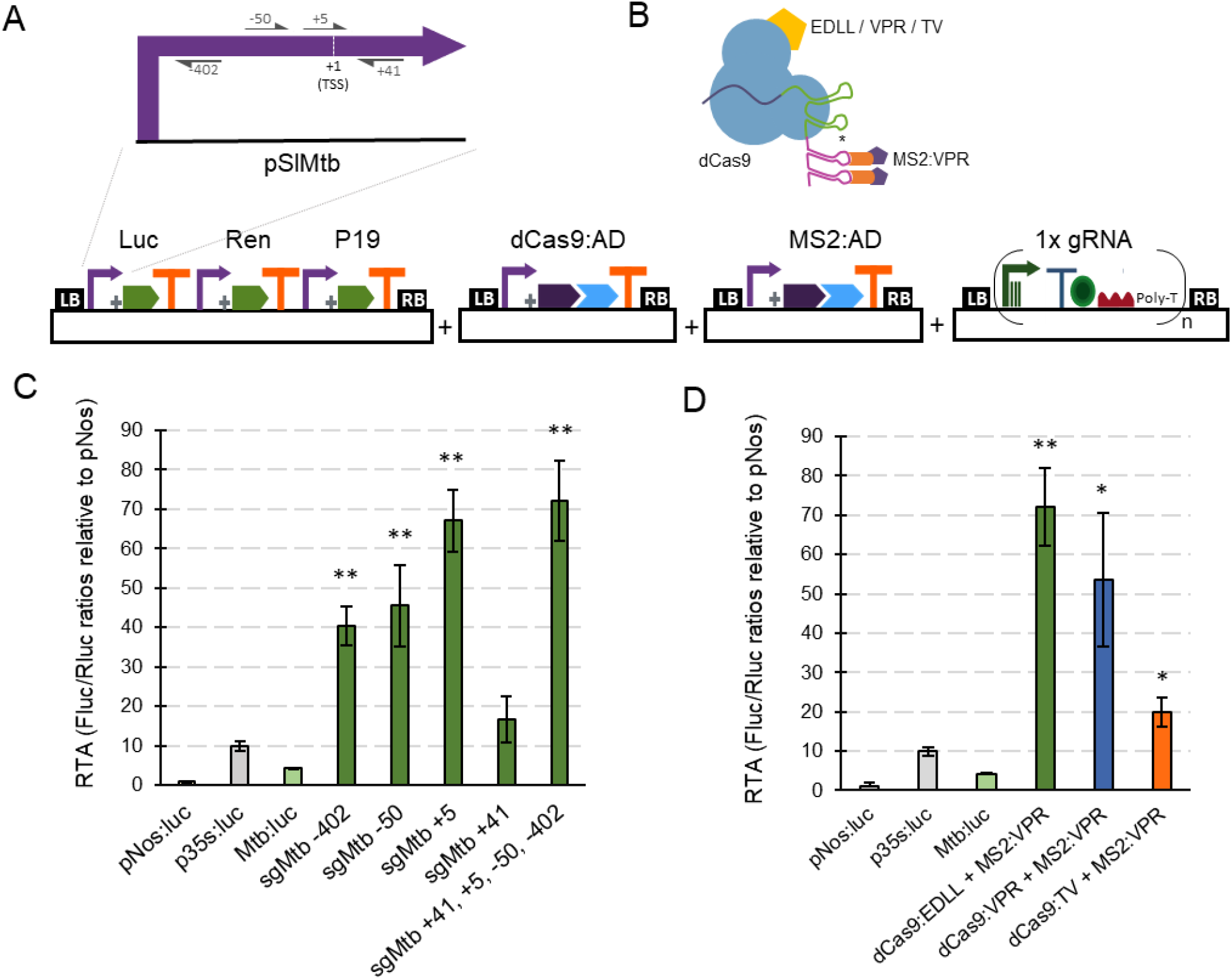
dCas9-based positive regulation in transient expression. A) Schematic representation of the GB plasmids co-infiltrated for evaluating dCas9-scRNA for activation of the *Solanum lycopersicum* Mtb promoter. B) Schematic representation of the dCas9-scRNA complex that includes the dCas9 fused to different activation domains and a gRNA with a 3’ extension on its scaffold that serves as anchoring site for the MS2:VPR protein. C) Relative transcriptional activities (RTA) of the tested gRNAs in combination with the dCas9:EDLL-MS2:VPR module and a luciferase reporter with the Mtb promoter. D) Comparison of relative transcriptional activities (RTA) obtained using different dCas9:AD (activation domain) versions. The data of bar charts represent the mean average of relative transcriptional activities (RTA) determined as Fluc/Rluc ratios of each sample normalized to Fluc/Rluc ratios of GB1116. The error bars indicate the standard deviations of all biological replicates (n=3). The statistical analyses were performed using unpaired *t* Test. Asterisks indicate significant differences with p35s:Luc with a \**P*-value < 0.05 and \*\**P*-value < 0.005.

Having established strong GB activator tools with dCas9-EV2.1, we next tested the ability of GB-made dCas12a-based constructs to repress promoter activity. The choice of dCas12a as scaffold for RDs responds to the convenience that both activators and repressors operate on different PAMs, therefore facilitating circuit design. Negative regulation is known to result from the activity of the repressor domain, but also from the steric interference of the transcription initiation and elongation complexes, a factor with strong positional dependence. Therefore, to optimize repression strategies we targeted several positions at the pNos promoter, and also inside the luciferase coding sequence (positions +201 and +380 in reference to the ATG) (Fig. 5A). From those, crRNAs targeting the pNos at -113, -33 and +26 resulted in significant repression rates of 42%, 41% and 53%, respectively (Fig. 5B). Notably, co-delivery of the three best crRNAs in a single T-DNA, either using single polycistronic TU or multiple TU approaches, resulted in maximum repression rates 58-67% respectively (Fig. 5B).

**Figure 5.**
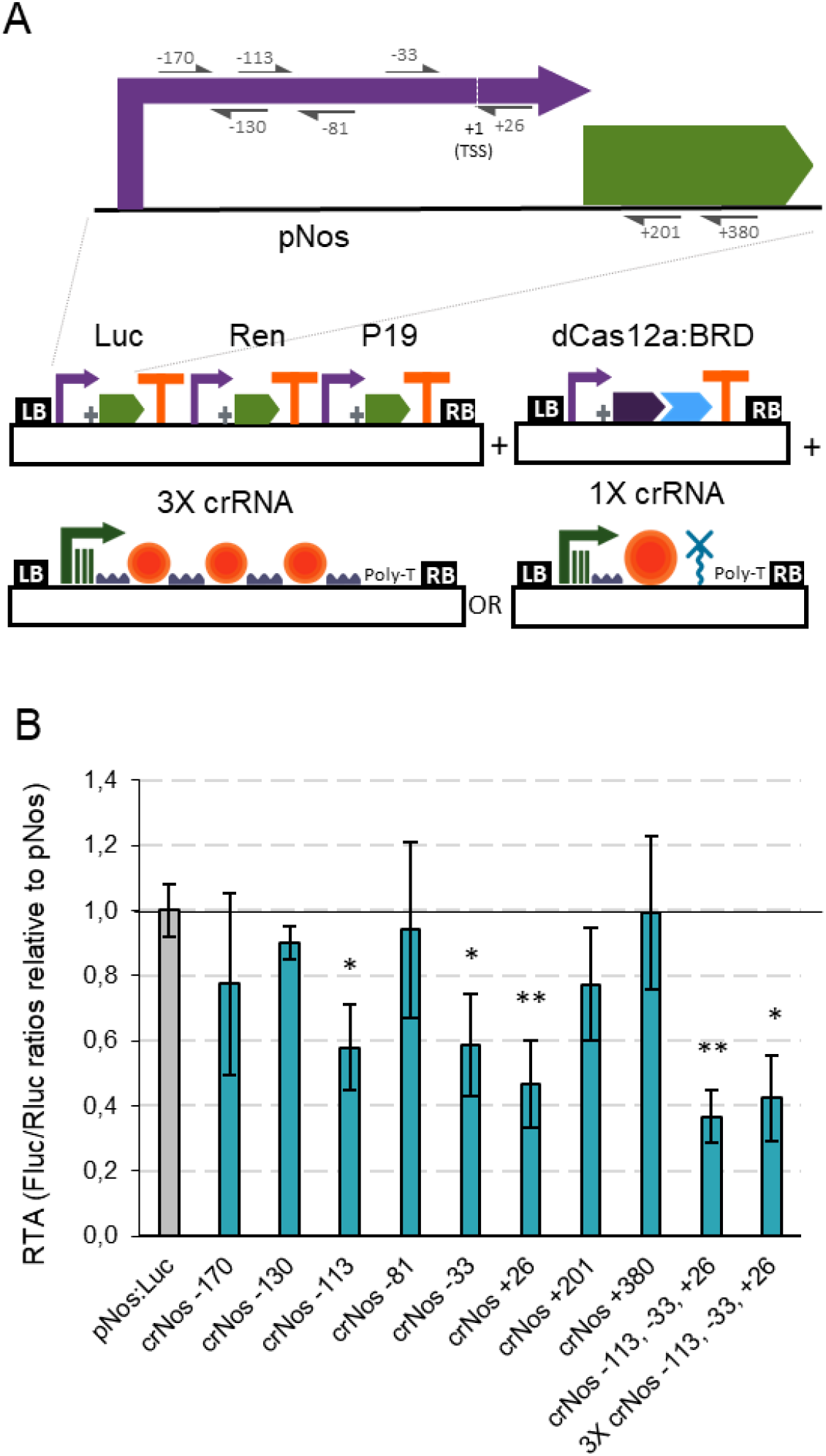
dCas12a-based negative regulation in transient expression. A) Schematic representation of the GB plasmids co-infiltrated for evaluating dCas12a as a tool for negative regulation of the nopaline synthase promoter. B) Relative transcriptional activities (RTA) of the tested gRNAs in combination with the dCas12a:BRD TU and a luciferase reporter with the nos promoter. The data of bar charts represent the mean average of relative transcriptional activities (RTA) determined as Fluc/Rluc ratios of each sample normalized to Fluc/Rluc ratios of GB1116. The error bars indicate the standard deviations of all biological replicates (n=3). The statistical analyses were performed using unpaired *t* Test. Asterisks indicate significant differences with pNos:Luc with a \**P*-value < 0.05 and \*\**P*-value < 0.005.

The opposite strategies, involving the use of GB-made dCas12a-based transcriptional activators and dCas9-based repressors, were also validated targeting several positions of the pNos promoter and the Firefly luciferase coding sequence. Maximum activation levels with dCas12a:VP64 were approximately two times the basal transcriptional levels of the pNos (Fig. S4) and repression levels of pNos with dCas9:BRD were slightly lower than those observed with dCas12a:BRD, even when targeting the Firefly luciferase coding sequence (Fig. S4).

## DISCUSSION

Virtually all areas of engineering owe part of their exponential growth to the implementation of standard rules and the development of functionally-documented components collections. Plant genetic engineering is unlikely to be an exception (Patron, 2020). GoldenBraid is a well-established multigene engineering platform using standard Type IIS RE-based assembly rules, and hosting currently circa 700 public physical phytobricks and >13.000 user-exclusive virtual gene elements, including basic DNA parts, transcriptional units and multigene modules, all of them assembled using the GB software. The GB platform has provided solutions to challenging multigene engineering endeavours as the engineering of the twelve-gene cholesterol pathway in Arabidopsis (Sonawane et al., 2016), the three genes betalain pathway in tomato (Polturak et al., 2016) or the engineering of a memory switch in *N. benthamiana* (Bernabé-Orts et al., 2020). In the advent of the plant genome engineering era, this fourth version of GB aims to offer extended capabilities to facilitate genome editing, taking advantage of the easy cloning and its software integration. GB4.0 includes a set of functionally tested plasmids for CRISPR editing, transcriptional activation and repression both with Cas9 and Cas12a as well as software-assisted cloning procedures for single and polycistronic gRNA assemblies. Other improvements in GB4.0 include the introduction of two new set of destination vectors in the cloning scheme with the ability to co-transform in the same Agrobacterium strain (Pasin et al., 2017). We have also agreed with the vSBOL community the use of new glyphs to represent different regions of the CDS, exon-intron boundaries and non-coding RNAs. Consequently, GB conforms to the most extended synthetic biology standard for DNA parts representation and vSBOL was accommodated to include visual representation for eukaryotic specific DNA parts (Baig et al., 2020).

CRISPR multiplexing offers unprecedented breeding potential, but construct assembly is often challenging and cumbersome. The GB4.0 genome edition offers a highly flexible solution for Cas9 and Cas12a multiplexing, providing all elements required for building any gRNA tandem comprising from 1 to 6 components and overcoming some of the limitations of previously described cloning strategies such as PCR dependency (Zuckermann et al., 2018). Cas9 has no ability to process its own gRNA tandem arrays, although recent studies in viral vectors seem to provide exceptions to this general rule (Uranga et al., 2020). Therefore, processable spacers need to be included in the array, so they can be rightly processed and trimmed into single functional units. Among the different spacer strategies described, we chose to incorporate the tRNA spacer method described by Xie et al. (Xie et al., 2015) to the GoldenBraid multiplexing pipeline. This method relies in endogenous plant RNase P and RNase Z required to process the tRNAs flanking each protospacer-scaffold unit. Other strategies involve removable spacers include the Csy4 RNase (requiring the supply in trans of the nuclease), or a self-cleavable hammerhead ribozyme (Gao and Zhao, 2014; Čermák et al., 2017). Comparative studies have found little differences in efficiency among the different systems available (Tang et al., 2019). Since the Csy4 system requires the supply in trans of the nuclease, imposing additional cargo to the multigene constructs, we favoured the use of the tRNA method, which on the other hand is best-suited for modular cloning. In the case of Cas12a, where gRNA processing activity of Cas12a itself is well established, the GB-proposed multiplexing strategy relayed on the direct chemical synthesis of the basic arrays, which are later assembled using regular GB binary-iterative cloning. Following GB cloning schema, tandems can be combined indefinitely using the iterative cloning pipeline. In an effort to reduce barriers in creating multiplex constructs, we developed an extremely simplify webtool, CRISPR for Dummies, which enables minimally trained users to create up to 6X gRNAs ready-to-transform editing construct in a pCAMBIA backbone following a step-by-step guided protocol. We are confident this and other tools in the website will pave the way for many labs to undergo challenging editing projects which would have been technically out of reach without GB tools in place. All the Cas9 and Cas12a, single and multiple gene editing systems available in the GB4.0 have been functionally validated, and in the particular case of Cas9 multiplexing, we have performed a stress test to the system by building and testing up to 17 gRNAs in the same construct. The physical linkage of Cas9 and all 17 gRNAs not only ensures multiple targeting but also simplifies the segregation of transgene-free T1 plants. The integration of DsRed positive selection marker greatly simplifies the segregation process (Aliaga-Franco et al., 2019).

Editing efficiency in plant is highly target-dependent, and despite many efforts to develop predictive algorithms (Haeussler et al., 2016), most researchers opt to follow an empirical approach, either performing prior efficiency tests *in vivo* using transient transformation methods (e.g. plant protoplasts), or by using force brute approaches i.e. targeting each gene at several positions. Targeting efficiency is known to be strongly affected by the gRNA sequence itself, with the chromatin accessibility of the target having probably also an influence (Naim et al., 2020). All the gRNAs employed in our stable transformation experiments were selected using the ‘Rule Set 2 scoring’ algorithm (Doench et al., 2016), showing all except one score values above 45. However, by introducing complex gRNA tandem arrays, additional considerations need to be made concerning the position of each gRNA in the array. The scale of the editing experiments described in this work, involving 28 gRNAs, >1000 editing events in a total of 13 HRs, 22 T0 and 55 T1 plants, allowed us to detect influences of the gRNA position in the editing efficiency. Our data shows that all positions are functional, however the last position in the array tends to produce higher efficiency levels, despite the predicted on-target score. A possible explanation for this bias is that the last position is flanked by a single tRNA, whereas the remaining positions need the release of both 5’and 3’ flanking tRNAS to produce a functional gRNA. If the nuclear supply of RNase P and RNase Z is a limiting factor, this could explain the observed differences. Alternatively, an increased stability of the 3’end of the RNA due to the presence of the polyA in the last position could also explain this preponderance. Whatever the mechanisms, this bias should be considered in the design of multiplexing constructs, as it might be advisable for instance to locate the most important guides (e.g. those targeting a larger number of orthologues) in the last position. In other occasions, it may be advisable to favour smaller tandems to maximize the number of last-position gRNAs, reaching a balance between multiplexing force and guide efficiency. Despite the observed bias, it becomes clear that multiplex strategies provide unprecedented power to breeding programs, especially in polyploid species, as exemplified with the MPO approach shown here, which yielded a 6X full knock out Cas9-free plant in a T1 generation. Interestingly, besides obtaining full-knock outs, multiplexing has the strength to generate a new type of “concentrated” genetic variability for large gene families. This custom variability has the ability to unveil phenotypes that remained hidden to natural diversity, especially in polyploid species, due to high functional redundancy. In the case of SPL genes, the T1 plants show strong phenotypes involving flowering time, branching and leaf juvenility among others that will be reported elsewhere, which could not have been generated without a concentrated multiplexing approach.

Finally, the assembly of functional modules to dCas has resulted in a variety of new powerful tools as programmable transcriptional regulators, base editors or prime editing elements among others (Shrestha et al., 2018; Mishra et al., 2020; Lin et al., 2020). We have focused in enriching the GB collection with elements that facilitate programmable transcriptional regulation. Whereas the successful activation on inducible genes has been earlier reported, here we show that CRISPR activators can also over-induce a strong promoter, boosting its activity above the level of CaMV35S promoter. The high expression levels obtained with dCas9EV2.1-activated pSlMTB suggest that this strategy could be exploited to boost yields of recombinant proteins and/or metabolites in biofactory and/or metabolic engineering approaches. In combination with Cas12a programmable repression in a multiplexing-enabled context, the new tools provide the ability to divert and tune endogenous metabolic pathways, channelling towards the production of metabolites of interest. Furthermore, the use of different Cas enzymes for positive and negative regulators also facilitates the design of genetic switches. However, in our hands negative regulation is still only partially efficient, and new improvements will be required to achieve stronger repression rates that allows the introduction of Boolean logic approximations for the design of genetic circuits (McCarty et al., 2020).

As in previous versions of the GoldenBraid system, version 4.0 provides a platform from which users can develop their own sub-collections. In the past, other groups have created independently GB extensions for plastids, mitochondria, yeast, geminiviral vectors, filamentous fungi, mammalian cells or amoebae (Vafaee et al., 2014; Pérez-González et al., 2017; Dahan-Meir et al., 2018; Hernanz-Koers et al., 2018; Sarrion-Perdigones et al., 2019; Kundert et al., 2020). Some extensions have been incorporated to the GB web, such as FungalBraid for filamentous fungi (Vazquez-Vilar et al., 2020), whereas others are internally developed by individual labs. In the current level of development, we have focused in nuclease-based SSN1 editing and transcriptional regulation. Obvious extensions like base editors or epigenetic editors can be incorporated with no effort within the GB4.0 frame, since they only require the addition of new level 0 parts to the collection, some of which have been already developed by users and collaborators. However, at this stage we only share in the public domain elements which have been functionally validated in our laboratory. Other extensions involving introduction of new functional elements in the gRNA, such as prime editing, would require additional software tools for full integration into the GB database. We are confident that the development of multipurpose platforms as GB4.0 is the way to go for turning plant biotechnology into a fast-advancing truly engineering discipline, and this can make a difference in the accessibility of many labs to genome editing and other related technologies.

## METHODS

### Guide RNA assembly on Level 0 and Level 1 plasmids

All Cas9 gRNAs used in this work with editing and repression purposes were assembled on Level 0 plasmids following the procedure described on Fig. S1 for Cas9 multiplexing. For the assembly of guide RNAs on level 0, two partially complementary primers designed at https://gbcloning.upv.es/do/crispr/multi_cas9_gRNA_domesticator_1 using as input the sequences listed on Table S1, were included in a BsmBI restriction–ligation reaction together with the pUPD2 and the corresponding level -1 tRNA-scaffold plasmid depending on the desired position of each target on the level 1 assembly. Level -1 tRNA-scaffold plasmid selection and assembly planning was done at https://gbcloning.upv.es/do/crispr/multi_cas9_gRNA_domesticator_2/. Cas9 gRNAs used for activation were assembled as Level 1 constructs following the single gRNA strategy (Fig. S1) using partially complementary primers designed at https://gbcloning.upv.es/do/crispr/Single_Cas9_gRNA_Domesticator. All level 0 and Level 1 gRNA constructs were validated by RE-analysis, analyzed by sequencing and confirmed correct.

Cas12a crRNAs were assembled using the procedures described in Fig. S1 for Cas12a single and multiplexing crRNAs. For Cas12a single crRNAs two partially complementary primers were designed at https://gbcloning.upv.es/do/crispr/Single_Cas12a_gRNA_Domesticator using as input the sequences listed on Table S1 and included in a BsaI restriction-ligation reaction together with GB1443, GB1444 and pDGB3alpha as a destination vector. For the assembly of the Cas12a 3X crRNA, a synthetic DNA fragment was designed at https://gbcloning.upv.es/tools/cas12multiplexing_domestication/, purchased from GenScript and cloned as a Level 0 DNA part prior to its assembly with GB1443 in a Level 1 plasmid. All Level 0 and Level 1 crRNA plasmids were validated by RE-analysis, analyzed by sequencing and confirmed correct.

### Cloning in α and Ω-level destination vectors

Multipartite BsaI restriction–ligation reactions from level 0 parts and binary BsaI or BsmBI restriction–ligation reactions were performed as described in (Vazquez-Vilar et al., 2020) to obtain all the level ≥1 assemblies. A list with all the TUs and modules used in this study is provided on Table S2. All level ≥1 plasmids were validated by restriction enzyme (RE) analysis. The sequences of all level ≥1 constructs can be found entering their IDs (displayed at Table S2) at https://gbcloning.upv.es/search/features/.

### Plant material

The *Nicotiana benthamiana* laboratory strain was used for all transient expression assays. *N. tabacum* cv. K326 was used for stable transformation either with *A. tumefaciens* or with *A. rhizogenes*. For the *A. rhizogenes* experiment, wild type tobacco plants were used. For the *A. tumefaciens* experiment, a line previously developed in the laboratory using CRISPR/Cas9 with homozygous mutations in six SPL genes and bi-allelic heterozygous mutations in one SPL gene (SPL157-5 T1) was used as genetic background for retransformation after T-DNA segregation. Edited genes and mutations of the SPL157-5 T1 line are: Nitab4.5_0003348g0050.1 (213delC), Nitab4.5_0007487g0020.1 (204delCA), Nitab4.5_0002219g0060.1 (213insA), Nitab4.5_0000638g0040.1 (559delGGACACAA / 565delAA), Nitab4.5_0001752g0040.1 (353delACAAC), Nitab4.5_0003572g0010.1 (289insC), Nitab4.5_0000016g0300.1 (535insT). In between brackets number indicates the position of the mutation in reference to the ATG, ‘del’ states for deletion, ‘ins’ states for insertion. It should be noticed that GB2714, include one gRNA (gSPL3.1) targeting Nitab4.5_0003348g0050.1 at a position different to that mutated on plant SPL157-5. Therefore, editing efficiencies of this gRNA targeting this specific gene are also included in Fig. 3C.

### *N. benthamiana* transient expression

For transient expression, plasmids were introduced into *Agrobacterium tumefaciens* strain GV3101 by electroporation. *N. benthamiana* plants were grown for 5 to 6 weeks before agroinfiltration in a growing chamber at 24 °C (light)/20 °C (darkness) with a 16-h-light/8-h-dark photoperiod. Agroinfiltration was carried out with overnight-grown bacterial cultures. The cultures were pelleted and resuspended on agroinfiltration solution (10 mM MES, pH 5.6, 10 mM MgCl2, and 200 μM acetosyringone) to an optical density of 0.1 at 600 nm. After incubation for 2 h at room temperature on a horizontal rolling mixer, the bacterial suspensions described for each experiment were mixed in equal volumes. The silencing suppressor P19 was included in all the assays; in the same T-DNA for the transcriptional regulation experiments and co-delivered in an independent T-DNA for the targeted mutagenesis assays. Agroinfiltration was carried out through the abaxial surface of the three youngest leaves of each plant with a 1 ml needle-free syringe.

### *N. tabacum* cv. K326 stable transformation

*Agrobacterium tumefaciens* LBA4404 harboring plasmid GB2484 was used to transform *Nicotiana tabacum* cultivar K326 following a standard protocol (Horsch, R., Fry, J., Hoffman, N., Eichholtz, D., Rogers, S.A. and Fraley, 1985). Briefly, fully expanded leaves of T1 157-5 plants were sterilized with 5% commercial bleach for 10 minutes followed by four consecutive washing steps with sterile demi water. Leaf discs (d= 0.8 cm) were cut with a cork borer and incubated overnight in co-culture plates (4.9 g/L MS supplemented with vitamins (Duchefa, https://www.duchefa-biochemie.com/), 3% sucrose (Sigma-Aldrich, https://www.sigmaaldrich.com/), 0.8% Phytoagar (Duchefa), 1 mg/L BAP (Sigma-Aldrich), 0.1 mg/L NAA (Sigma-Aldrich), pH=5.7). Leaf discs were incubated for 15 minutes with a culture of *A. tumefaciens* LBA4404 harboring plasmid GB2714 (OD600=0.3). Then, the discs were returned to the co-cultivation plates and incubated for 2 days in darkness. Next, discs were transferred to selection medium (4.9 g/L MS supplemented with vitamins (Duchefa), 3% sucrose (Sigma-Aldrich), 0.8% Phytoagar (Duchefa), 1 mg/L BAP (Sigma-Aldrich), 0.1 mg/L NAA (Sigma-Aldrich), 500 mg/L carbenicillin, 100 mg/L kanamycin, pH=5.7). Discs were transferred to fresh medium every seven days until shoots appeared (4-6 weeks). Shoots were cut and transferred to rooting medium (4.9 g/L MS supplemented with vitamins (Duchefa), 3% sucrose (Sigma-Aldrich), 0.8% Phytoagar (Duchefa), 500 mg/L carbenicillin, 100 mg/L kanamycin, pH=5.7) until roots appeared.

*Agrobacterium rhizogenes* strain ATCC® 15834 (Kuzma et al., 1995) carrying the plasmid GB2484 was used for transformation of WT tobacco leaves to obtain hairy roots. After leaf sterilization following the protocol described above, leaf discs were incubated overnight in co-cultivation plates without BAP and without NAA. Leaf discs were incubated for 30 minutes with shaking in a culture of *A. rhizogenes* previously grown overnight in YEB medium with 200 uM acetosyringone, pelleted and resuspended in MS (2.45 g/L MS supplemented with vitamins (Duchefa), 3% sucrose (Sigma-Aldrich), pH=5.7) to OD600=0.6. Next, the leaf discs were transferred to co-cultivation plates without BAP and without NAA and incubated for 2 days in darkness. Then, the discs were transferred to selection plates (4.9 g/L MS supplemented with vitamins (Duchefa), 3% sucrose (Sigma-Aldrich), 0.8% Phytoagar (Duchefa), 500 mg/L carbenicillin, 100 mg/L kanamycin, pH=5.7). Discs were transferred to fresh medium every seven days until roots appeared (3-4 weeks). Hairy roots were isolated from the leaf disc and transferred to pots with 50 ml of liquid selection medium (4.9 g/L MS supplemented with vitamins (Duchefa), 3% sucrose (Sigma-Aldrich), 500 mg/L carbenicillin, 100 mg/L kanamycin, pH=5.7). Roots were grown for 1 month until enough material for genomic DNA extraction was obtained. Growing conditions were in all steps 16 h light/ 8 h dark, 25°C, 60-70% humidity, 250 µmol m-2 s-1 photons.

### Genomic DNA extraction and editing efficiency evaluation

150 mg of either leaf material from T0 tobacco stable transformants or from 5 dpi *N. benthamiana* leaves or root material from tobacco hairy roots was used for genomic DNA extraction. Genomic DNA was extracted with the CTAB (cetyl trimethylammonium bromide) method (Murray and Thompson, 1980). The genomic regions flanking the nuclease target sites were PCR amplified using MyTaq™ DNA Polymerase (Bioline, https://www.bioline.com/) and primers listed on Table S3. The PCR amplicons were confirmed on a 1% agarose gel electrophoresis and purified with ExoSAP-IT™ PCR Product Cleanup Reagent (ThermoFisher Scientific, https://www.thermofisher.com) following the manufacturer’s indications prior to Sanger sequencing. Chromatograms of Cas9-edited genomic DNA were analyzed using Inference of CRISPR Edits (ICE) v2 tool from Synthego (https://ice.synthego.com/) and chromatograms of Cas12a-edited genomic DNA were analyzed with TIDE (http://shinyapps.datacurators.nl/tide/). All analysis were manually curated. When two or more gRNAs targeting the same gene show a 50% of editing efficiency it is not possible to determine from the chromatogram analysis if one or both alleles are mutated. On these cases estimations were done in the most conservative way and heterozygous mutations were assumed.

### Sampling and Luciferase assays

Samples of leaves coinfiltrated with the pNos or pMtb reporter construct (REP, GB1116 or GB1399, respectively), different activator/repressor TUs or modules (GB1830, GB1190, GB1826, GB2047, GB1668, GB1665, GB1172) and the individual or polycistronic gRNAs targeting either the pNos or the pMtb (Table S1 and S2) were collected at 4 days post infiltration. For the determination of the luciferase/renilla activity one disc per leaf (d = 0.5 cm) was excised, homogenized and extracted with 180 µl of ‘Passive Lysis Buffer’, followed by 15 min of centrifugation (14,000 ×g) at 4 °C. Fluc and Rluc activities were determined following the Dual-Glo® Luciferase Assay System (Promega, https://www.promega.es/) manufacturer’s protocol with minor modifications: 10 µl of plant extract, 40 µl of LARII and 40 µl of Stop&Glo Reagent were used. Measurements were made using a GloMax 96 Microplate Luminometer (Promega) with a 2-s delay and a 10-s measurement. Fluc/Rluc ratios were determined as the mean value of three samples coming from three independent agroinfiltrated leaves of the same plant and were normalized to the Fluc/Rluc ratio obtained for a reference sample including the REP (GB1116). GB1119, a p35S reporter construct was included as upper limit reference in all experiments.

## AUTHOR CONTRIBUTIONS

M.V., S.S., J.B. and D.O. designed the experiments. M.V., S.S., J.B., J.S., B.S., A.R., C.P. M.A. conducted the experiments. V.G. developed the software tools. A.F. designed the website. M.V. and D.O. drafted the manuscript. M.V., A.F., A.G. and D.O. discussed and revised the manuscript. All the authors read and approved the final manuscript.

## ACKNOWLEDGEMENTS

The authors would like to thank the ‘IBMCP sequencing service’ staff, Eugenio Grau and Ana Marín, for their excellent assistance with Sanger sequencing. This work has been funded by EU Horizon 2020 Project Newcotiana Grant 760331 and BIO2016-78601-R Plan Nacional I+D, Spanish Ministry of Economy and Competitiveness. Vazquez-Vilar, M. is recipient of APOSTD/2020/096 (Generalitat Valenciana and Fondo Social Europeo post-doctoral grant). Bernabé-Orts, J.M. and Selma, S. are recipients of FPI fellowships. De Paola, C. is recipient of a Santiago Grisolía fellowship (Generalitat Valenciana).

The authors declare no conflict of interest.

## REFERENCES

Aliaga-Franco, N., Zhang, C., Presa, S., Srivastava, A. K., Granell, A., Alabadí, D., Sadanandom, A., Blázquez, M. A., and Minguet, E. G. (2019). Identification of Transgene-Free CRISPR-Edited Plants of Rice, Tomato, and Arabidopsis by Monitoring DsRED Fluorescence in Dry Seeds. Front Plant Sci 10.

Baig, H., Fontanarrosa, P., Kulkarni, V., McLaughlin, J., Vaidyanathan, P., Bartley, B., Bhatia, S., Bhakta, S., Bissell, M., Clancy, K., et al. (2020). Synthetic biology open language visual (SBOL visual) version 2.2. Journal of Integrative Bioinformatics 17.

Bernabé-Orts, J. M., Casas-Rodrigo, I., Minguet, E. G., Landolfi, V., Garcia-Carpintero, V., Gianoglio, S., Vázquez-Vilar, M., Granell, A., and Orzaez, D. (2019). Assessment of Cas12a-mediated gene editing efficiency in plants. Plant Biotechnology Journal Advance Access published 2019, DOI:10.1111/pbi.13113.

Bernabé-Orts, J. M., Quijano-Rubio, A., Vazquez-Vilar, M., Mancheño-Bonillo, J., Moles-Casas, V., Selma, S., Gianoglio, S., Granell, A., and Orzaez, D. (2020). A memory switch for plant synthetic biology based on the phage ϕC31 integration system. Nucleic Acids Res 48:3379–3394.

Čermák, T., Curtin, S. J., Gil-Humanes, J., Čegan, R., Kono, T. J. Y., Konečná, E., Belanto, J. J., Starker, C. G., Mathre, J. W., Greenstein, R. L., et al. (2017). A multipurpose toolkit to enable advanced genome engineering in plants. Plant Cell 29:1196–1217.

Concordet, J.-P., and Haeussler, M. (2018). CRISPOR: intuitive guide selection for CRISPR/Cas9 genome editing experiments and screens. Nucleic Acids Res 46:W242–W245.

Dahan-Meir, T., Filler-Hayut, S., Melamed-Bessudo, C., Bocobza, S., Czosnek, H., Aharoni, A., and Levy, A. A. (2018). Efficient in planta gene targeting in tomato using geminiviral replicons and the CRISPR/Cas9 system. The Plant Journal 95:5–16.

Doench, J. G., Fusi, N., Sullender, M., Hegde, M., Vaimberg, E. W., Donovan, K. F., Smith, I., Tothova, Z., Wilen, C., Orchard, R., et al. (2016). Optimized sgRNA design to maximize activity and minimize off-target effects of CRISPR-Cas9. Nat Biotechnol 34:184–191.

Engler, C., Gruetzner, R., Kandzia, R., and Marillonnet, S. (2009). Golden gate shuffling: A one-pot DNA shuffling method based on type ils restriction enzymes. PLoS ONE Advance Access published 2009, DOI:10.1371/journal.pone.0005553.

Gao, Y., and Zhao, Y. (2014). Self-processing of ribozyme-flanked RNAs into guide RNAs in vitro and in vivo for CRISPR-mediated genome editing. Journal of Integrative Plant Biology 56:343–349.

Haeussler, M., Schönig, K., Eckert, H., Eschstruth, A., Mianné, J., Renaud, J.-B., Schneider-Maunoury, S., Shkumatava, A., Teboul, L., Kent, J., et al. (2016). Evaluation of off-target and on-target scoring algorithms and integration into the guide RNA selection tool CRISPOR. Genome Biology 17:148.

Hernanz-Koers, M., Gandía, M., Garrigues, S., Manzanares, P., Yenush, L., Orzaez, D., and Marcos, J. F. (2018). FungalBraid: A GoldenBraid-based modular cloning platform for the assembly and exchange of DNA elements tailored to fungal synthetic biology. Fungal Genetics and Biology 116:51–61.

Horsch, R., Fry, J., Hoffman, N., Eichholtz, D., Rogers, S.A. and Fraley, R. (1985). A Simple and General Method for Transferring Genes into Plants. Science Advance Access published 1985, DOI:10.1126/science.227.4691.1229.

Jansing, J., Sack, M., Augustine, S. M., Fischer, R., and Bortesi, L. (2019). CRISPR/Cas9-mediated knockout of six glycosyltransferase genes in Nicotiana benthamiana for the production of recombinant proteins lacking β-1,2-xylose and core α-1,3-fucose. Plant Biotechnology Journal 17:350–361.

Kang, B., Yun, J., Kim, S., Shin, Y., Ryu, J., Choi, M., Woo, J. W., and Kim, J. (2018). Precision genome engineering through adenine base editing in plants. Nature Plants 4.

Kim, H. K., Song, M., Lee, J., Menon, A. V., Jung, S., Kang, Y.-M., Choi, J. W., Woo, E., Koh, H. C., Nam, J.-W., et al. (2017). In vivo high-throughput profiling of CRISPR–Cpf1 activity. Nature Methods 14:153–159.

Konermann, S., Brigham, M. D., Trevino, A. E., Joung, J., Abudayyeh, O. O., Barcena, C., Hsu, P. D., Habib, N., Gootenberg, J. S., Nishimasu, H., et al. (2015). Genome-scale transcriptional activation by an engineered CRISPR-Cas9 complex. Nature Advance Access published 2015, DOI:10.1038/nature14136.

Kundert, P., Sarrion-Perdigones, A., Gonzalez, Y., Katoh-Kurasawa, M., Hirose, S., Lehmann, P., Venken, K. J. T., and Shaulsky, G. (2020). A GoldenBraid cloning system for synthetic biology in social amoebae. Nucleic Acids Res. 48:4139–4146.

Kuzma, J., Nemecek-Marshall, M., Pollock, W. H., and Fall, R. (1995). Bacteria produce the volatile hydrocarbon isoprene. Current Microbiology Advance Access published 1995, DOI:10.1007/BF00294190.

Kwon, C. T., Heo, J., Lemmon, Z. H., Capua, Y., Hutton, S. F., Van Eck, J., Park, S. J., and Lippman, Z. B. (2020). Rapid customization of Solanaceae fruit crops for urban agriculture. Nature Biotechnology 38:182–188.

Lin, Q., Zong, Y., Xue, C., Wang, S., Jin, S., Zhu, Z., Wang, Y., Anzalone, A. V., Raguram, A., Doman, J. L., et al. (2020). Prime genome editing in rice and wheat. Nat Biotechnol 38:582–585.

McCarty, N. S., Graham, A. E., Studená, L., and Ledesma-Amaro, R. (2020). Multiplexed CRISPR technologies for gene editing and transcriptional regulation. Nature Communications 11:1–13.

Mishra, R., Joshi, R. K., and Zhao, K. (2020). Base editing in crops: current advances, limitations and future implications. Plant Biotechnol. J. 18:20–31.

Naim, F., Shand, K., Hayashi, S., O’Brien, M., McGree, J., Johnson, A. A. T., Dugdale, B., and Waterhouse, P. M. (2020). Are the current gRNA ranking prediction algorithms useful for genome editing in plants? PLoS ONE 15:1–12.

Papikian, A., Liu, W., Gallego-Bartolomé, J., and Jacobsen, S. E. (2019). Site-specific manipulation of Arabidopsis loci using CRISPR-Cas9 SunTag systems. Nature Communications Advance Access published 2019, DOI:10.1038/s41467-019-08736-7.

Pasin, F., Bedoya, L. C., Bernabé-Orts, J. M., Gallo, A., Simón-Mateo, C., Orzaez, D., and García, J. A. (2017).https://pubs.acs.org/doi/pdf/10.1021/acssynbio.6b00354 Accessed September 23, 2020.

Patron, N. J. (2020). Beyond natural: synthetic expansions of botanical form and function. New Phytol 227:295–310.

Patron, N. J., Orzaez, D., Marillonnet, S., Warzecha, H., Matthewman, C., Youles, M., Raitskin, O., Leveau, A., Farré, G., Rogers, C., et al. (2015). Standards for plant synthetic biology: A common syntax for exchange of DNA parts. New Phytologist Advance Access published 2015, DOI:10.1111/nph.13532.

Pérez-González, A., Kniewel, R., Veldhuizen, M., Verma, H. K., Navarro-Rodríguez, M., Rubio, L. M., and Caro, E. (2017). Adaptation of the GoldenBraid modular cloning system and creation of a toolkit for the expression of heterologous proteins in yeast mitochondria. BMC Biotechnol 17:80–80.

Polturak, G., Breitel, D., Grossman, N., Sarrion-Perdigones, A., Weithorn, E., Pliner, M., Orzaez, D., Granell, A., Rogachev, I., and Aharoni, A. (2016). Elucidation of the first committed step in betalain biosynthesis enables the heterologous engineering of betalain pigments in plants. New Phytologist 210:269–283.

Sánchez-León, S., Gil-Humanes, J., Ozuna, C. V., Giménez, M. J., Sousa, C., Voytas, D. F., and Barro, F. (2018). Low-gluten, nontransgenic wheat engineered with CRISPR/Cas9. Plant Biotechnology Journal 16:902–910.

Sarrion-Perdigones, A., Falconi, E. E., Zandalinas, S. I., Juárez, P., Fernández-del-Carmen, A., Granell, A., and Orzaez, D. (2011). GoldenBraid: An iterative cloning system for standardized assembly of reusable genetic modules. PLoS ONE Advance Access published 2011, DOI:10.1371/journal.pone.0021622.

Sarrion-Perdigones, A., Vazquez-Vilar, M., Palací, J., Castelijns, B., Forment, J., Ziarsolo, P., Blanca, J., Granell, A., and Orzaez, D. (2013). Goldenbraid 2.0: A comprehensive DNA assembly framework for plant synthetic biology. Plant Physiology Advance Access published 2013, DOI:10.1104/pp.113.217661.

Sarrion-Perdigones, A., Chang, L., Gonzalez, Y., Gallego-Flores, T., Young, D. W., and Venken, K. J. T. (2019). Examining multiple cellular pathways at once using multiplex hextuple luciferase assaying. Nature communications 10:5710.

Selma, S., Bernabé-Orts, J. M., Vazquez-Vilar, M., Diego-Martin, B., Ajenjo, M., Garcia-Carpintero, V., Granell, A., and Orzaez, D. (2019). Strong gene activation in plants with genome-wide specificity using a new orthogonal CRISPR /Cas9-based programmable transcriptional activator. Plant Biotechnology Journal Advance Access published 2019, DOI:10.1111/pbi.13138.

Shimatani, Z., Kashojiya, S., Takayama, M., Terada, R., Arazoe, T., Ishii, H., Teramura, H., Yamamoto, T., Komatsu, H., Miura, K., et al. (2017). Targeted base editing in rice and tomato using a CRISPR-Cas9 cytidine deaminase fusion. Nature Publishing Group 35:441–443.

Shrestha, A., Khan, A., and Dey, N. (2018). cis–trans Engineering: Advances and Perspectives on Customized Transcriptional Regulation in Plants. Molecular Plant 11:886–898.

Sonawane, P. D., Pollier, J., Panda, S., Szymanski, J., Massalha, H., Yona, M., Unger, T., Malitsky, S., Arendt, P., Pauwels, L., et al. (2016). Plant cholesterol biosynthetic pathway overlaps with phytosterol metabolism. Nature Plants 3:1–13.

Tanenbaum, M. E., Gilbert, L. A., Qi, L. S., Weissman, J. S., and Vale, R. D. (2014). A protein-tagging system for signal amplification in gene expression and fluorescence imaging. Cell Advance Access published 2014, DOI:10.1016/j.cell.2014.09.039.

Tang, X., Ren, Q., Yang, L., Bao, Y., Zhong, Z., He, Y., Liu, S., Qi, C., Liu, B., Wang, Y., et al. (2019). Single transcript unit CRISPR 2.0 systems for robust Cas9 and Cas12a mediated plant genome editing. Plant Biotechnology Journal 17:1431–1445.

Uranga, M., Aragonés, V., Selma, S., Vázquez-Vilar, M., Orzáez, D., and Daròs, J.-A. (2020). Efficient Cas9 multiplex editing using unspaced gRNA arrays engineering in a Potato virus X vector. bioRxiv Advance Access published June 26, 2020, DOI:10.1101/2020.06.25.170977.

Vafaee, Y., Staniek, A., Mancheno-Solano, M., and Warzecha, H. (2014). A modular cloning toolbox for the generation of chloroplast transformation vectors. PLoS One 9:e110222.

Vazquez-Vilar, M., Quijano-Rubio, A., Fernandez-Del-Carmen, A., Sarrion-Perdigones, A., Ochoa-Fernandez, R., Ziarsolo, P., Blanca, J., Granell, A., and Orzaez, D. (2017). GB3.0: a platform for plant bio-design that connects functional DNA elements with associated biological data. Nucleic acids research Advance Access published 2017, DOI:10.1093/nar/gkw1326.

Vazquez-Vilar, M., Gandía, M., García-Carpintero, V., Marqués, E., Sarrion-Perdigones, A., Yenush, L., Polaina, J., Manzanares, P., Marcos, J. F., and Orzaez, D. (2020). Multigene Engineering by GoldenBraid Cloning: From Plants to Filamentous Fungi and Beyond. Current Protocols in Molecular Biology Advance Access published 2020, DOI:10.1002/cpmb.116.

Weber, E., Engler, C., Gruetzner, R., Werner, S., and Marillonnet, S. (2011). A modular cloning system for standardized assembly of multigene constructs. PLoS ONE Advance Access published 2011, DOI:10.1371/journal.pone.0016765.

Xie, K., Minkenberg, B., and Yang, Y. (2015). Boosting CRISPR/Cas9 multiplex editing capability with the endogenous tRNA-processing system. Proceedings of the National Academy of Sciences of the United States of America 112:3570–3575.

Zhang, Y., Pribil, M., Palmgren, M., and Gao, C. (2020). A CRISPR way for accelerating improvement of food crops. Nature Food Advance Access published 2020, DOI:10.1038/s43016-020-0051-8.

Zheng, M., Zhang, L., Tang, M., Liu, J., Liu, H., Yang, H., Fan, S., Terzaghi, W., Wang, H., and Hua, W. (2020). Knockout of two BnaMAX1 homologs by CRISPR/Cas9-targeted mutagenesis improves plant architecture and increases yield in rapeseed (Brassica napus L.) Plant Biotechnol J 18:644–654.

Zhu, H., and Liang, C. (2019). CRISPR-DT: designing gRNAs for the CRISPR-Cpf1 system with improved target efficiency and specificity. Bioinformatics 35:2783–2789.

Zong, Y., Wang, Y., Li, C., Zhang, R., Chen, K., Ran, Y., Qiu, J., Wang, D., and Gao, C. (2017). Precise base editing in rice, wheat and maize with a Cas9-cytidine deaminase fusion. Nature Publishing Group 35:438–440.

Zsögön, A., Čermák, T., Naves, E. R., Notini, M. M., Edel, K. H., Weinl, S., Freschi, L., Voytas, D. F., Kudla, J., and Peres, L. E. P. (2018). De novo domestication of wild tomato using genome editing. Nature Biotechnology 36:1211–1216.

Zuckermann, M., Hlevnjak, M., Yazdanparast, H., Zapatka, M., Jones, D. T. W., Lichter, P., and Gronych, J. (2018). A novel cloning strategy for one-step assembly of multiplex CRISPR vectors. Sci Rep 8.

