## Supplementary material for "Edition of complex gene families in tobacco with GoldenBraid 4.0, a multipurpose web-based platform for plant genome engineering": Table S1, Table S2, Table S1 and S2, Table S3, Table S4, Fig. S1, Fig. S2, Fig. S3, Fig. S4

### Supplementary information

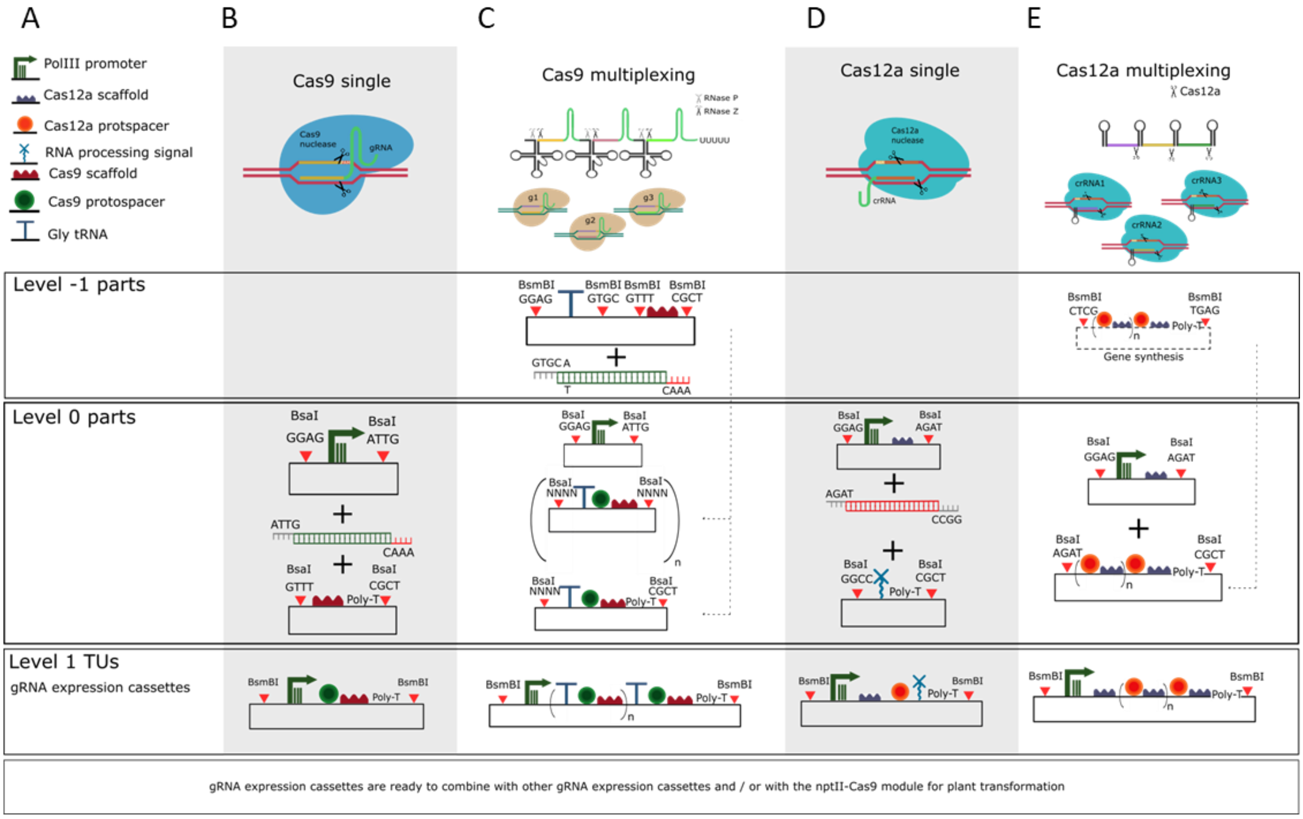

**Supplementary Figure 1. Cas9 and Cas12a single and multiplexing guide RNA expression cassettes cloning strategies with GoldenBraid.** A) Visual SBOL glyphs for the Level 0 parts involved in the gRNAs expression cassettes assembly. B) Cas9 single gRNAs are assembled as Level 1 constructs with a PolIII promoter, hybridized primers including the protospacer sequence and the corresponding overhangs, and the Cas9 scaffold. C) Cas9 multiplexing gRNAs assembly involves the assembly of Level 0 tRNA-protospacer-scaffold units using hybridized primers including the protospacer sequences and the corresponding overhangs, and the corresponding Level -1 vectors. Level 0 tRNA-protospacer-scaffold units are assembled together and with a PolII promoter to create a Level 1 polycistronic gRNA expression cassette. D) Cas12a single gRNAs are assembled as Level 1 constructs with a PolIII promoter including the Cas12a scaffold, hybridized primers including the protospacer sequences and the corresponding overhangs and a 3’ RNA processing signal. E) Cas12a multiplexing gRNAs assembly involves the synthesis of (protospacer-scaffold)$_n$ and the subsequent cloning of the synthetic fragment as a Level 0 part for further assembly with a PolIII promoter including the Cas12a scaffold in Level 1.

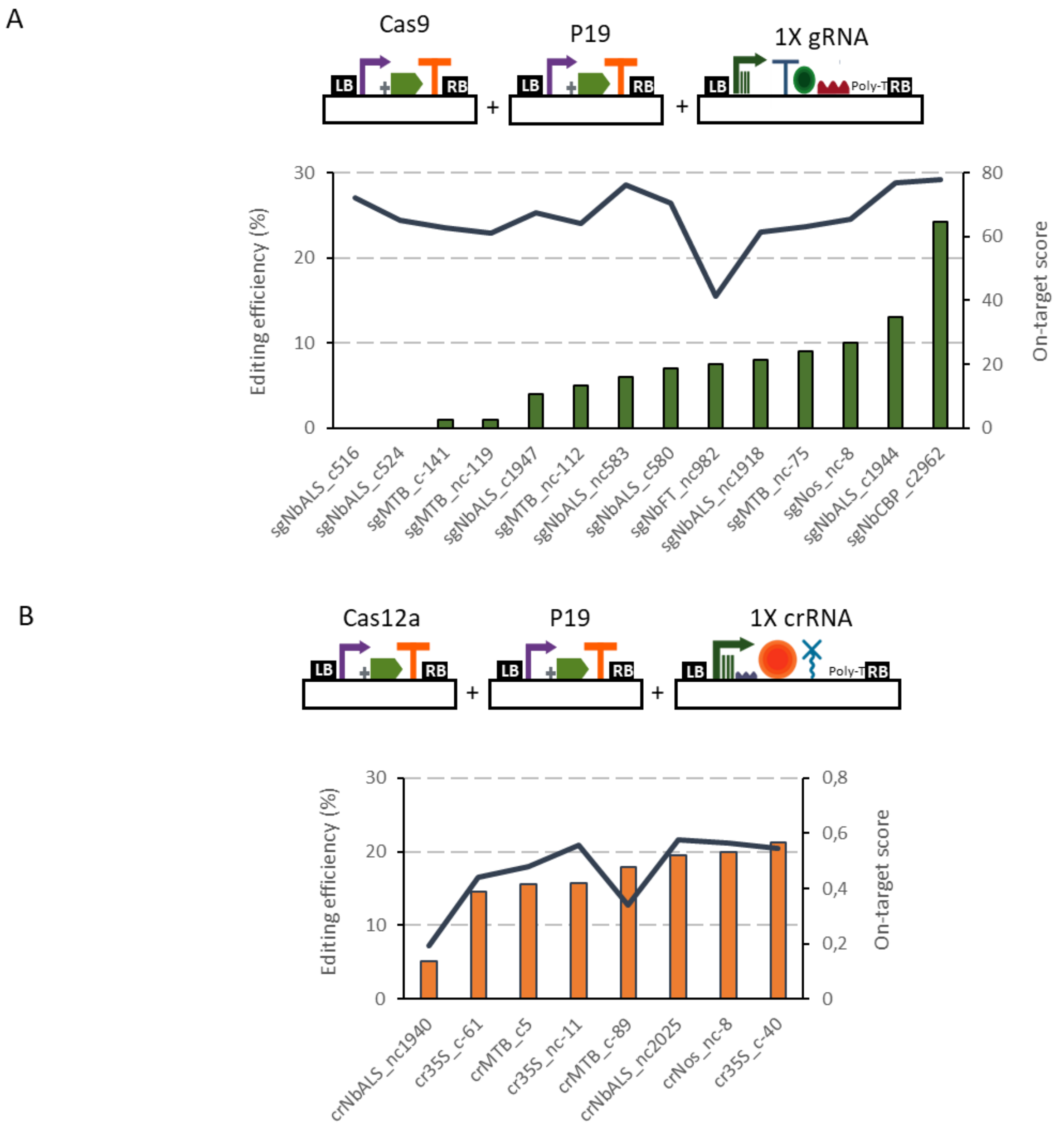

**Supplementary Figure 2. Correlation of Cas9 and Cas12a guideRNAs predicted on-target scores and editing efficiencies tested in *N. benthamiana* transient expression.** A) Schematic representation of the plasmids co-infiltrated in this experiment (top) and Cas9 guide RNAs editing efficiencies (left axis, bars, determined with ICE) and their corresponding on-target score (right axis, line) determined with the 'Rule Set2 scoring' (bottom). B) Schematic representation of the plasmids co-infiltrated in this experiment (top) and Cas12a guide RNAs editing efficiencies (left axis, bars, determined with TIDE) and their corresponding on-target score (right axis, line) determined with CINDEL (bottom).

A

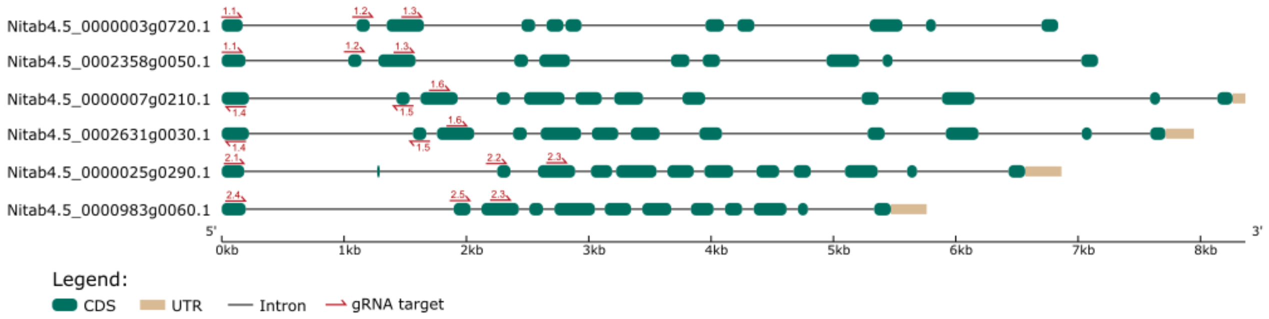

B

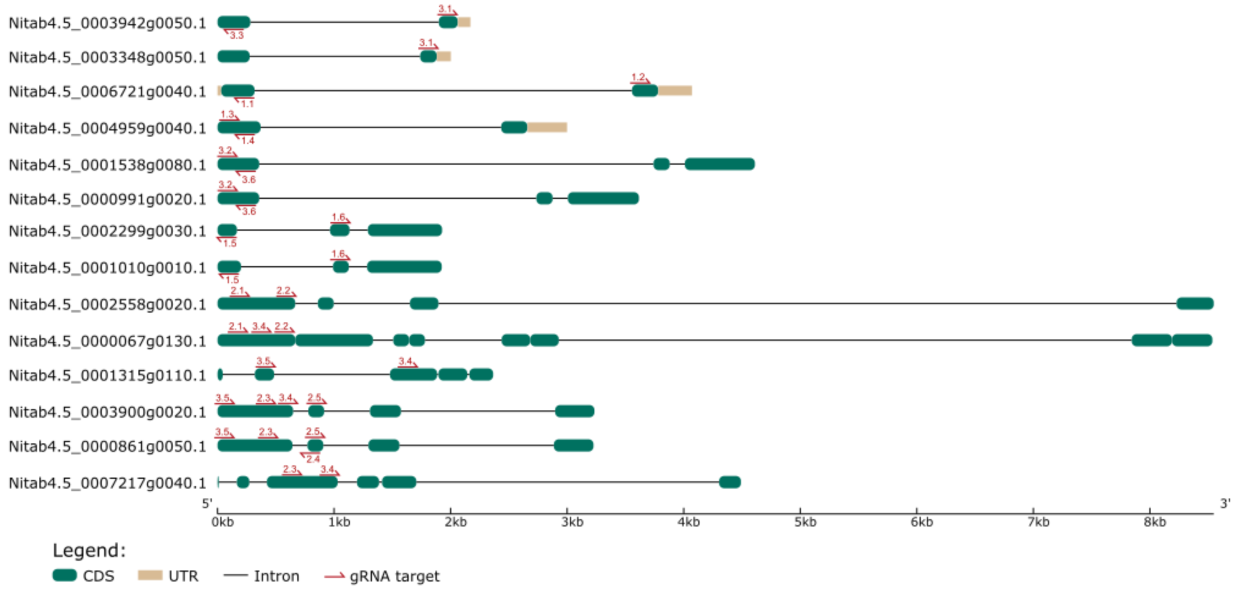

**Supplementary Figure 3. Schematic representation of the exon-intron gene structure for the targeted genes.** A) Exon-intron representation of the targeted MPO genes indicating the approximate positions targeted by each gRNA in GB2484. B) Exon-intron representation of the targeted SPL genes indicating the approximate positions targeted by each gRNA in GB2714.

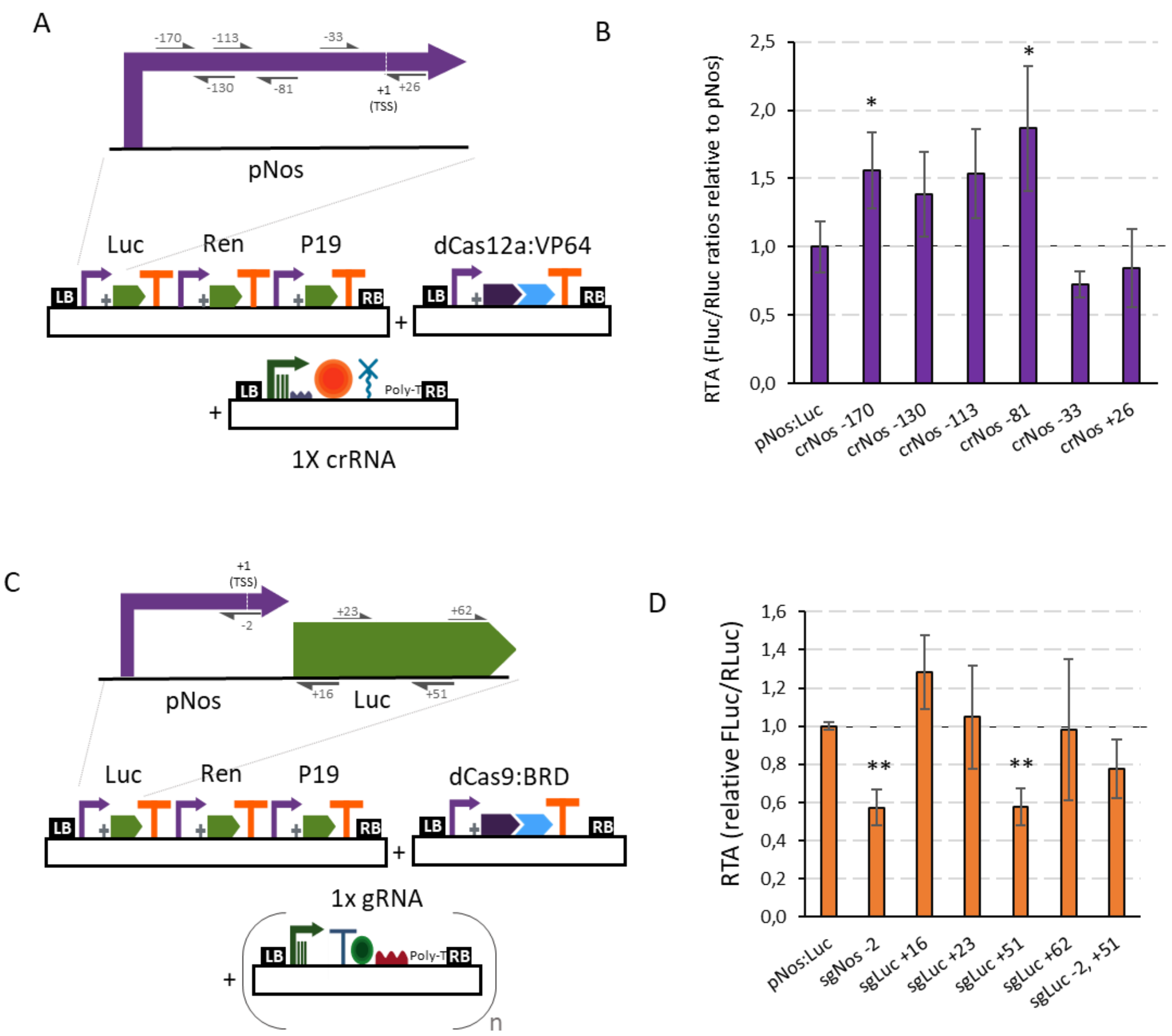

**Supplementary Figure 4. dCas12a-based transcriptional upregulation and dCas9-based transcriptional downregulation in transient expression in *N. benthamiana*.** A) Schematic representation of the GB plasmids co-infiltrated for evaluating dCas12a:VP64 for activation of the nopaline synthase promoter. B) Relative transcriptional activities (RTA) of the tested crRNAs in combination with the dCas12a:VP64 TU and a luciferase reporter with the nos promoter. C) Schematic representation of the GB plasmids co-infiltrated for evaluating dCas9 as a tool for negative regulation of the nopaline synthase promoter. D) Relative transcriptional activities (RTA) of the tested gRNAs in combination with the dCas9:BRD TU and a luciferase reporter with the nos promoter. The data of bar charts represent the mean average of relative transcriptional activities (RTA) determined as Fluc/Rluc ratios of each sample normalized to Fluc/Rluc ratios of GB1116. The error bars indicate the standard deviations of all biological replicates (n=3). The statistical analyses were performed using unpaired *t* Test. Asterisks indicate significant differences with pNos:Luc with a **P*-value < 0.05 and ***P*-value < 0.005.

**Table S1. List of protospacer sequences and targeted genes used in this study.** EDI stands for editing, ACT for activation and REP for repression. *indicates scores estimated with 23 nt genomic sequences corresponding to the crRNA plus additional nucleotides up to 23 nts (CINDEL score is only available for crRNAs of length ≥ 23 nts). gRNA targeted positions were determined as distance of the Cas9/Cas12a cutting site to the ATG for coding sequences and to the TSS for promoter regions. "c" indicates that the gRNA is designed on the coding strand and "nc" that gRNA is designed on the non-coding strand.

| gRNA name | Targeted gene(s) and targeted position | Protospacer sequence | PAM | On-target score[1,2] | Cas | Purpose |
|---|---|---|---|---|---|---|
| gSPL1.1 | Nitab4.5_0006721g0040.1nc260 | CGACGGCTTGAGACTTTGCA | TGG | 64 | Cas9 | EDI |
| gSPL1.2 | Nitab4.5_0006721g0040.1c363 | AATCGAAAAGGAGTTGCAGG | CGG | 70 | Cas9 | EDI |
| gSPL1.3 | Nitab4.5_0004959g0040.1c142 | AAGGGTCAAGTACTTCAATG | AGG | 72 | Cas9 | EDI |
| gSPL1.4 | Nitab4.5_0004959g0040.1nc631 | CGACGGCTTGAGCCTTTGCA | TGG | 60 | Cas9 | EDI |
| gSPL1.5 | Nitab4.5_0002299g0030.1nc135/ Nitab4.5_0001010g0010.1nc123 | GGACCTCACAAACTTTATGG | CGG | 70 | Cas9 | EDI |
| gSPL1.6 | Nitab4.5_0002299g0030.1c274/ Nitab4.5_0001010g0010.1c277 | ATGGACATAACAGGCGTCGA | AGG | 61 | Cas9 | EDI |
| gSPL2.1 | Nitab4.5_0000067g0130.1c419/ Nitab4.5_0002558g0020.1c419 | CGAACAGTTGAAAGCAACAG | TGG | 69 | Cas9 | EDI |
| gSPL2.2 | Nitab4.5_0000067g0130.1c640/ Nitab4.5_0002558g0020.1c640 | TCATCATAGGAGGTGCCGAG | CGG | 74 | Cas9 | EDI |
| gSPL2.3 | Nitab4.5_0000861g0050.1c334/ Nitab4.5_0003900g0020.1c340 / Nitab4.5_0007217g0040.1c466 | TAGAAATTCACCCCCCATGG | AGG | 66 | Cas9 | EDI |
| gSPL2.4 | Nitab4.5_0000861g0050.1nc692 | TGTGATCAGAGAGCCTCCTG | CGG | 66 | Cas9 | EDI |
| gSPL2.5 | Nitab4.5_0003900g0020.1c757 | GGAAACCATCCAGTTCAACT | CGG | 66 | Cas9 | EDI |
| gSPL3.1 | Nitab4.5_0003348g0050.1_c408 / Nitab4.5_0003942g0050.1_c414 | TTAAAGGGGCATCAGTCCAA | TGG | 63 | Cas9 | EDI |
| gSPL3.2 | Nitab4.5_0000991g0020.1_c122 / Nitab4.5_0001538g0080.1_c122 | ATCTACTTTGAAAATGTGGG | TGG | 65 | Cas9 | EDI |
| gSPL3.3 | Nitab4.5_0003348g0050.1_nc212 / Nitab4.5_0003942g0050.1_nc224 | GAAATTCACAGACCTTGTGG | CGG | 73 | Cas9 | EDI |
| gSPL3.4 | Nitab4.5_0001315g0110.1c406/ Nitab4.5_0007217g0040.1c723 / Nitab4.5_0003900g0020.1c598 / Nitab4.5_0000067g0130.1c616 | CAGTCATTCCAAATGCCCAA | AGG | 69 | Cas9 | EDI |
| gSPL3.5 | Nitab4.5_0001315g0110.1c192 | AGATGGAGTGGAATGCAAAG | TGG | 67 | Cas9 | EDI |
| gSPL3.6 | Nitab4.5_0000991g0020.1_nc216 / Nitab4.5_0001538g0080.1_nc216 | CCTTCAACTTGACACCTAGG | TGG | 70 | Cas9 | EDI |
| gMPO1.1 | Nitab4.5_0000003g0720.1c112 / Nitab4.5_0002358g0050.1c136 | CGATCAGCAGAAACAAACGC | CGG | 82 | Cas9 | EDI |
| gMPO1.2 | Nitab4.5_0000003g0720.1c277 / Nitab4.5_0002358g0050.1c301 | AGCTGCCGGTGAAACACCCG | AGG | 21 | Cas9 | EDI |
| gMPO1.3 | Nitab4.5_0000003g0720.1c536 / Nitab4.5_0002358g0050.1c560 | GCTCGAGGTGGACATCACAA | GGG | 95 | Cas9 | EDI |
| gMPO1.4 | Nitab4.5_0000007g0210.1nc88 / Nitab4.5_0002631g0030.1nc88 | AATTTCAGCTACGGTTCCGG | CGG | 67 | Cas9 | EDI |
| gMPO1.5 | Nitab4.5_0000007g0210.1nc295 / Nitab4.5_0002631g0030.1nc295 | CCTGACAGTTGCAACAGCCA | CGG | 67 | Cas9 | EDI |
| gMPO1.6 | Nitab4.5_0000007g0210.1c587 / Nitab4.5_0002631g0030.1c587 | ACTCGAGGAGGCCACCACCG | CGG | 69 | Cas9 | EDI |
| gMPO2.1 | Nitab4.5_0000025g0290.1nc80 | CGGCAGCTGTCGCCTCACGA | CGG | 47 | Cas9 | EDI |

| | | | | | | |
|---|---|---|---|---|---|---|
| gMPO2.2 | Nitab4.5_0000025g0290.1c223 | AGGTATACAAATCATGACAA | GGG | 49 | Cas9 | EDI |
| gMPO2.3 | Nitab4.5_0000025g0290.1c572 / Nitab4.5_0000983g0060.1c590 | GCTCGAGGTGGACATCACAG | GGG | 91 | Cas9 | EDI |
| gMPO2.4 | Nitab4.5_0000983g0060.1nc92 | TGGCCGCTGTTGCCTCACGA | CGG | 57 | Cas9 | EDI |
| gMPO2.5 | Nitab4.5_0000983g0060.1c241 | AGGTATCCAAATCATGACAA | GGG | 54 | Cas9 | EDI |
| sgNbALS_c516 | Niben101Scf05032_c516 | GGCGCCACTAATCTCGTCAG | CGG | 72 | Cas9 | EDI |
| sgNbALS_c524 | Niben101Scf05032_c524 | TAATCTCGTCAGCGGCCTCG | CGG | 65 | Cas9 | EDI |
| sgNbALS_c580 | Niben101Scf05032_c580 | TAACCGGTCAAGTGCCACGT | AGG | 70 | Cas9 | EDI |
| sgNbALS_nc583 | Niben101Scf05032_nc583 | TCAGTACCGATCATCCTACG | TGG | 76 | Cas9 | EDI |
| sgNbALS_nc1918 | Niben101Scf05032_nc1918 | GGTAGAACATGTTCCTGATG | AGG | 61 | Cas9 | EDI |
| sgNbALS_c1944 | Niben101Scf05032_c1944 | GTTCTACCTATGATTCCCAG | TGG | 77 | Cas9 | EDI |
| sgNbALS_c1947 | Niben101Scf05032_c1947 | CTACCTATGATTCCCAGTGG | CGG | 68 | Cas9 | EDI |
| crNbALS_c1940 | Niben101Scf05032_c1940 | AAAGCTCCGCCACTGGGAATCAT | TTTG | 19 | Cas12a | EDI |
| crNbALS_c2025 | Niben101Scf05032_c2025 | AGATAATACAAGGTCTAGAACTA | TTTC | 58 | Cas12a | EDI |
| sgNos _c-8 | Nopaline synthase promoter_c-8 | ATTCACTCTCAACTCGATCG | AGG | 65 | Cas9 | EDI |
| crNos:Luc_nc-8 | Nopaline synthase promoter_nc-8 | GCGTCTTCCATTGCCTCGATCGA | TTTG | 57 | Cas12a | EDI |
| sgSlMtb_c-141 | Solyc09g010800_c-141 | ATACGATCACGACACGTGTA | CGG | 63 | Cas9 | EDI |
| sgSlMtb_nc-119 | Solyc09g010800_nc-119 | TATAGTGAAACGAAGGTACA | TGG | 61 | Cas9 | EDI |
| sgSlMtb_nc-112 | Solyc09g010800_nc-112 | GTCTATTTATAGTGAAACGA | AGG | 64 | Cas9 | EDI |
| sgSlMtb_nc-75 | Solyc09g010800_nc-75 | GGTGTAGAAATGAGTGATTG | AGG | 63 | Cas9 | EDI |
| crMtb_c-89 | Solyc09g010800_c-89 | ACTATAAATAGACACTCATGCTT | TTTC | 0.34 | Cas12a | EDI |
| crMtb:Luc_c5 | Firefly Luciferase_c5 | AGACCAATCAATCAATGGAAGAC | TTTC | 0.48 | Cas12a | EDI |
| cr35s_c-62 | CaMV35s_c-62 | GAGAGGACTCCGGTATTTTTACA | TTTG | 0.44 | Cas12a | EDI |
| cr35s_c-41 | CaMV35s_c-41 | CAACAATTACCACAACAAAACAA | TTTA | 0.54 | Cas12a | EDI |
| cr35s:Ren_nc-11 | CaMV35s:Renilla_nc-11 | GAAGTCATTTCGACTAGAATAGT | TTTC | 0.56 | Cas12a | EDI |
| sgCBP_c487 | Niben101Scf02459g00024.1_c487 | GAGGACAAACTACATCCAGG | TGG | 78 | Cas9 | EDI |
| sgFT_nc62 | Niben101Scf01519g10008.1_nc62 | GGCCAATAGATCTTGTAAAA | GGG | 45 | Cas9 | EDI |
| crFT_c83 | Niben101Scf01519g10008.1_c83 | CAAGATCTATTGGCCTAAGAG | TTTA | 0.23 | Cas12a | EDI |
| sgSlMTB_nc41 | Solyc09g010800_nc41 (-155 in reference to ATG) | GTACACGTGTCGTGATCGTA | TGG | 55 | Cas9 | ACT |
| sgSlMTB_c5 | Solyc09g010800_c5 (-191 in reference to ATG) | GATGAAATTAGGATCATGTA | TGG | 52 | Cas9 | ACT |
| sgSlMTB_c-50 | Solyc09g010800_c-50 (-246 in reference to ATG) | GTCTAGAACATACGTACGAA | TGG | 59 | Cas9 | ACT |
| sgSlMTB_nc-402 | Solyc09g010800_nc-402 (-598 in reference to ATG) | GATGTAGCATATGAGATGAT | AGG | 60 | Cas9 | ACT |
| crLuc_nc380 | Firefly Luciferase_nc380 | CAACCCCTTTTTGGAAACGA | TTTG | 0.18 | Cas12a | REP |
| crLuc_nc201 | Firefly Luciferase_nc201 | TATTCAGCCCATATCGTTTC | TTTG | 0.11 | Cas12a | REP |

| | | | | | | |
|---|---|---|---|---|---|---|
| crNos_nc26 | Nopaline synthase promoter_nc26 (-9 in reference to ATG) | GCGTCTTCCATTGCCTCGAT | TTTG | 0.24 | Cas12a | ACT/REP |
| crNos_nc-33 | Nopaline synthase promoter_nc-33 (-67 in reference to the ATG) | TTGTCAAAAATGCTCCACTG | TTTC | 0.08 | Cas12a | ACT/REP |
| crNos_nc-81 | Nopaline synthase promoter_nc-81 (-115 in reference to the ATG) | CTAGCTGATAGTGACCTTAG | TTTG | 0.18 | Cas12a | ACT/REP |
| crNos_nc-113 | Nopaline synthase promoter_nc-113 (-147 in reference to the ATG) | TGACGTATGTGCTTAGCTCA | TTTC | 0.13 | Cas12a | ACT/REP |
| crNos_nc-130 | Nopaline synthase promoter_nc-130 (-164 in reference to the ATG) | ATGAGCTAAGCACATACGTC | TTTA | 0.33 | Cas12a | ACT/REP |
| crNos_nc-170 | Nopaline synthase promoter_nc-170 (-204 in reference to the ATG) | GAACTGACAGAACCGCAACG | TTTG | 0.25 | Cas12a | ACT/REP |
| sgNos_nc-2 | Nopaline synthase promoter_nc-2 (-36 in reference to the ATG) | AGTGAATATGAGACTCTAAT | TGG | 61 | Cas9 | REP |
| sgLuc_nc16 | Firefly Luciferase_nc16 (-35 in reference to ATG) | CGGGCCTTTCTTTATGTTTT | TGG | 10 | Cas9 | REP |
| sgLuc_c23 | Firefly Luciferase_c23 | GACGCCAAAAACATAAAGAA | AGG | 52 | Cas9 | REP |
| sgLuc_nc51 | Firefly Luciferase_nc51 | CCAGCGGTTCCATCTTCCAG | CGG | 66 | Cas9 | REP |
| sgLuc_c62 | Firefly Luciferase_c62 | CCGCTGGAAGATGGAACCGC | TGG | 47 | Cas9 | REP |

[1] Doench, J. G., Fusi, N., Sullender, M., Hegde, M., Vaimberg, E. W., Donovan, K. F., Smith, I., Tothova, Z., Wilen, C., Orchard, R., et al. (2016). Optimized sgRNA design to maximize activity and minimize off-target effects of CRISPR-Cas9. *Nat Biotechnol* 34:184–191.

[2]Kim, H. K., Song, M., Lee, J., Menon, A. V., Jung, S., Kang, Y.-M., Choi, J. W., Woo, E., Koh, H. C., Nam, J.-W., et al. (2017). In vivo high-throughput profiling of CRISPR–Cpf1 activity. *Nature Methods* 14:153–159.

**Table S2. List of plasmids generated during this work.** All sequences and further plasmid's information can be searched at https://gbcloning.upv.es/search/features/ with the GB IDs.

| GB ID | Plasmid name |
|---|---|
| GB2714 | pEGB3o1_nptII-Cas9-DsRed-6xsgNtSPL1-5xsgNtSPL2-6xsgNtSPL3 (gSPL1.1-1.6 + gSPL2.1-2.5 + gSPL3.1-3.6) |
| GB2712 | pEGB3a1_nptII-Cas-DsRed-6xsgNtSPL1-5xsgNtSPL2 (gSPL1.1-1.6 + gSPL2.1-2.5) |
| GB2708 | pEGB3a2_6xsgNtSPL3 (gSPL3.1-gSPL3.6) |
| GB2235 | pEGB3omega1 tNos:nptII:PNos-35s:Cas9:tNos-35s:DsRed:tNos |
| GB2709 | pEGB3o2_6xsgNtSPL1-5xsgNtSPL2 (gSPL1.1-1.6 + gSPL2.1-2.5) |
| GB2706 | pEGB3a1_6xsgNtSPL1 (gSPL1.1-1.6) |
| GB2707 | pEGB3a2_5xsgNtSPL2 (gSPL2.1-2.5) |
| GB2626 | pUPD2_sgNtSPL1.1 [E1] |
| GB2627 | pUPD2_sgNtSPL1.2 [E2] |
| GB2628 | pUPD2_sgNtSPL1.3 [E3] |
| GB2629 | pUPD2_sgNtSPL1.4 [E4] |
| GB2630 | pUPD2_sgNtSPL1.5 [En-1] |
| GB2631 | pUPD2_sgNtSPL1.6 [EnC1] |
| GB2632 | pUPD2_sgNtSPL2.1 [E1] |
| GB2633 | pUPD2_sgNtSPL2.2 [E2] |
| GB2634 | pUPD2_sgNtSPL2.3 [E3] |
| GB2635 | pUPD2_sgNtSPL2.4 [E4-En-1] |
| GB2636 | pUPD2_sgNtSPL2.5 [EnC1] |
| GB2699 | pUPD2_sgNtSPL3.1 [E1] |
| GB2700 | pUPD2_sgNtSPL3.2 [E2] |
| GB2701 | pUPD2_sgNtSPL3.3 [E3] |
| GB2702 | pUPD2_sgNtSPL3.4 [E4] |
| GB2703 | pUPD2_sgNtSPL3.5 [En-1] |
| GB2704 | pUPD2_sgNtSPL3.6 [EnC1] |
| GB2484 | pEGB3a1_nptII-Cas9-DsRed-6XMPO1-5XMPO2 (gMPO1.1-1.6+gMPO2.1-2.5) |
| GB2460 | pEGB3o2_6XMPO1+5XMPO2 (gMPO1.1-1.6+gMPO2.1-2.5) |
| GB2458 | pEGB3a1_6XMPO1 (gMPO1.1-1.6) |
| GB2459 | pEGB3a2_5XMPO2 (gMPO2.1-2.5) |
| GB2447 | pUPD2-sgNtMPO1.1 [E1] |
| GB2448 | pUPD2-sgNtMPO1.2 [E2] |
| GB2449 | pUPD2-sgNtMPO1.3 [E3] |
| GB2450 | pUPD2-sgNtMPO1.4 [E4] |
| GB2451 | pUPD2-sgNtMPO1.5 [En-1] |
| GB2452 | pUPD2-sgNtMPO1.6 [EnC1] |
| GB2453 | pUPD2-sgNtMPO2.1 [E1] |
| GB2454 | pUPD2-sgNtMPO2.2 [E2] |
| GB2455 | pUPD2-sgNtMPO2.3 [E3] |
| GB2456 | pUPD2-sgNtMPO2.4 [E4-En-1] |
| GB2457 | pUPD2-sgNtMPO2.5 [EnC1] |
| GB2549 | pEGB3α2_U626:pre-tRNA:sgNbALS_c516:SpScaffold |
| GB2550 | pEGB 3α2_U626:pre-tRNA:sgNbALS_c524:SpScaffold |
| GB2551 | pEGB 3α2_U626:pre-tRNA:sgNbALS_c580:SpScaffold |
| GB2552 | pEGB 3α2_U626:pre-tRNA:sgNbALS_nc583:SpScaffold |
| GB3014 | pEGB3α2_U626:pre-tRNA:sgRNA NbALS 1918:SpScaffold |
| GB3015 | pEGB3α2_U626:pre-tRNA:sgRNA NbALS 1944: Scaffold |
| GB3016 | pEGB3α2_U626:pre-tRNA:sgRNA NbALS 1947: Scaffold |
| GB3017 | pEGB3α2_U626:crNbALS_c1940: HDV |
| GB3018 | pEGB3α2_U626:crNbALS_c2025: HDV |

| | |
|---|---|
| GB2553 | pEGB3α2_U626:pre-tRNA:sgpNos::Luc_c-8:Scaffold |
| GB2554 | pEGB3α2_U626:LbScaffold:crpNos::Luc_nc-8:HDV |
| GB2555 | pEGB3α2_U626:pre-tRNA:sgSlMtb_c-141:SpScaffold |
| GB2556 | pEGB3α2_U626:pre-tRNA:sgSlMtb _nc-119:SpScaffold |
| GB3583 | pEGB3α2_U626:pre-tRNA:sgSlMtb_nc-112:SpScaffold |
| GB3582 | pEGB3α2_U626:pre-tRNA:sgSlMtb _nc-75:SpScaffold |
| GB2557 | pEGB3α2_U626:LbScaffold:crSlMtb_c-89:HDV |
| GB2558 | pEGB3α2_U626:LbScaffold:crSlMtb::Luc_c5:HDV |
| GB2559 | pEGB3α2_U626:LbScaffold:cr35S_c-62:HDV |
| GB3030 | pEGB3α2_U626:LbScaffold:cr35S_c-41: HDV |
| GB2561 | pEGB3α2_U626:LbScaffold:cr35S::Ren_nc-11: HDV |
| GB1774 | pEGB3α2_U626:sgNbCBP_c487:SpScaffold |
| GB1773 | pEGB3α2_U626:sgNbFT_nc62:SpScaffold |
| GB3033 | pEGB3α2_U626:LbScaffold:crNbFT_c83: HDV |
| GB2541 | pUPD2_sgNbALS_c516 [E1-E2-E3-E4-En-1-EnC1] |
| GB2542 | pUPD2_ sgNbALS_c524 [E1-E2-E3-E4-En-1-EnC1] |
| GB2543 | pUPD2_ sgNbALS_c580 [E1-E2-E3-E4-En-1-EnC1] |
| GB2544 | pUPD2_ sgNbALS_nc583 [E1-E2-E3-E4-En-1-EnC1] |
| GB3002 | pUPD2_ sgNbALS_nc1918 [E1-E2-E3-E4-En-1-EnC1] |
| GB3003 | pUPD2_ sgNbALS_c1944 [E1-E2-E3-E4-En-1-EnC1] |
| GB3004 | pUPD2_ sgNbALS_c1947 [E1-E2-E3-E4-En-1-EnC1] |
| GB2545 | pUPD2_sgNos_c-8 [E1-E2-E3-E4-En-1-EnC1] |
| GB2546 | pUPD2_sgSlMtb_c-141 [E1-E2-E3-E4-En-1-EnC1] |
| GB2547 | pUPD2_sgSlMtb_nc-119 [E1-E2-E3-E4-En-1-EnC1] |
| GB2548 | pUPD2_sgSlMtb_nc-112 [E1-E2-E3-E4-En-1-EnC1] |
| GB3581 | pUPD2_sgSlMtb_nc-75 [E1-E2-E3-E4-En-1-EnC1] |
| GB1119 | pEGB 35s:Luc:TNos-SF-35s:Renilla:TNos-35s:P19:TNos-SF |
| GB1399 | pEGB3alpha2 MTB:luc:Tnos-SF-35S:Ren:Tnos-35s:P19:Tnos-SF |
| GB1830 | 35s:Ms2:VPR:tNos |
| GB2047 | 35s:dCas9:TV:tNos |
| GB1190 | 35s:dCas9:EDLL:tNos |
| GB1826 | 35s:dCas9:VPR:tNos |
| GB2045 | 3alpha1:U6-26:sgSlMtb_nc41_scf F6x2 |
| GB1801 | 3alpha1:U6-26:sgSlMtb_c5_scf F6x2 |
| GB2044 | 3alpha2:U6-26:sgSlMtb_c-50_scf F6x2 |
| GB1859 | 3alpha1:U6-26:sgSlMtb_nc-402_scf F6x2 |
| GB2070 | pEGB3alpha1_U6-26:sgSlMtb_nc-402:F6x2_U6-26:sgSlMtb_c-50:F6x2_U6-26:sgSlMtb_c5:F6x2_U6-26:sgSlMtb_nc41:F6x2 |
| GB1116 | pEGB pNos:Luciferase:TNos-SF-35S:Renilla:TNos-35S:P19:TNos-SF |
| GB1668 | pEGB 3alpha1 35S:dLbCas12a:BRD:T35S |
| GB1805 | pEGB3alpha2_ U626:LbScaffold:crLuc_nc380:HDV |
| GB1806 | pEGB3alpha2_ U626:LbScaffold:crLuc_nc201:HDV |
| GB1807 | pEGB3alpha2_ U626:LbScaffold:crpNos::Luc_nc26:HDV |
| GB1808 | pEGB3alpha2_ U626:LbScaffold:crpNos_nc-33:HDV |
| GB1809 | pEGB3alpha2_ U626:LbScaffold:crpNos_nc-81:HDV |
| GB1811 | pEGB3alpha2_ U626:LbScaffold:crpNos_nc-113:HDV |
| GB1812 | pEGB3alpha2_ U626:LbScaffold:crpNos_nc-130:HDV |
| GB1813 | pEGB3alpha2_ U626:LbScaffold:crpNos_nc-170:HDV |
| GB1879 | U6-26::LbScaffold:crNos_nc-113:LbScaffold:crNos-33:LbScaffold:crNos_nc26:LbDR |
| GB1172 | pEGB 35s:dCas9:BRD:tNOS |
| GB1197 | pEGB3alpha2 U626:sgNos_nc-2: SpScaffold |
| GB2647 | pEGB3alpha2 U626:sgNos_nc16: SpScaffold |
| GB2648 | pEGB3alpha2 U626:sgNos_c23: SpScaffold |
| GB2778 | pEGB3alpha2 U626:sgNos_nc51: SpScaffold |
| GB2779 | pEGB3alpha2 U626:sgNos_c62: SpScaffold |

| | |
|---|---|
| GB2642 | pUPD2 sgNos_nc16 [E1-E2-E3-E4-En-1-EnC1] |
| GB2643 | pUPD2 sgNos_c23 [E1-E2-E3-E4-En-1-EnC1] |
| GB2645 | pUPD2 sgNos_nc51 [E1-E2-E3-E4-En-1-EnC1] |
| GB2646 | pUPD2 sgNos_c62 [E1-E2-E3-E4-En-1-EnC1] |

**Table S3. List of primers used for amplification of the targeted sites.**

| Primer Name | Sequence (5’ → 3’) |
|---|---|
| MV19OCT01 SPL2299_g1.5_F3 | GTGTGATCGATGGAATCATCA |
| MV19SEP12 SPL2299_g1.5_R2 | ATCTTCCTAACTATGTGGCG |
| MV19AGO03 SPL1010_g1.5_F | GTGTGATCGATGGAATCATCC |
| MV19AGO04 SPL1010_g1.5_R | GTTTCCAGCCTTTATGAGAAC |
| MV19AGO05 SPL2299_g1.6_F | TGCTCTTGAGTCCTGTCC |
| MV19AGO06 SPL2299_g1.6_R | GTTGACTGCTAAATGACAGAAG |
| MV19AGO07 SPL1010_g1.6_F | ATGATGTGTGATTACTTAGCAC |
| MV19AGO08 SPL1010_g1.6_R | GTTGACTGCTAAATGACAGAAC |
| MV19AGO09 SPL0067_g2.1&2.2_F | GAGCTCAATATCAGCTTCTACC |
| MV19AGO10 SPL0067_g2.1&2.2_R | CTATACATCCAGAAGAACAAGTTGTT |
| MV19AGO11 SPL2558_g2.1&2.2_F | GAGAACTTGATCGCAGATCG |
| MV19AGO12 SPL2558_g2.1&2.2_R | GCTATACATCCAGAAGAACACG |
| MV19AGO13 SPL0861_g2.3&g2.4_F | GATTCGAACTTTCCTGTTGTTTC |
| MV19AGO14 SPL0861_g2.3&g2.4_R | CTTTAGCTGATGAAAGATCAAGGT |
| MV19AGO15 SPL3900_g2.3&g2.4_F | GTGTTCTTGCTCTTTGATTCA |
| MV19AGO16 SPL3900_g2.3&g2.4_R | CTTTAGCTGATGAAAGATCATGGC |
| MV19AGO17 SPL0861_g3.4&g2.5_F | GGCTCTGTTGAACCACTTG |
| MV19AGO18 SPL0861_g3.4&g2.5_R | CTGTCCATGTTAACCTATTCTC |
| MV19AGO19 SPL3900_g3.4&g2.5_F | GACATTTGAGACCATCGGC |
| MV19AGO20 SPL3900_g3.4&g2.5_R | GGTTGTTTAACCAGCAAAGAAAC |
| MV19AGO21 SPL7217_g3.5,g2.3&g3.4_F | GGAGATGCATTTTGTGGC |
| MV19AGO22 SPL7217_g3.5,g2.3&g3.4_R | CCAGTTGGGTGTGGTTGAGA |
| MV19AGO61 SPL1315_g3.5_F2 | CAACAGCTCTGTTTGATAAAGATCC |
| MV19AGO62 SPL1315_g3.5_R2 | CAAGTTCTCCATCTTCTTCAAC |
| MV19AGO25 SPL1315_g3.4_F | GGTGCTAAGATTTCATCCTTCG |
| MV19AGO26 SPL1315_g3.4_R | AACATGGATTCCTTGGCATC |
| MV19AGO27 SPL4959_g1.3&g1.4_F | CTTAGCCTATATCTTCCTATAGC |
| MV19AGO28 SPL4959_g1.3&g1.4_R | GTATTTTAGTAACTTAAAAATGTGCTG |
| MV19AGO29 SPL6721_g1.3&g1.1_F | CACTTAGCCTATATCTTCCTAAC |
| MV19AGO30 SPL6721_g1.3&g1.4_R | CAACATGCTAAAGAATGCATC |
| MV19AGO31 SPL6721_g1.2_F | TTCATACACACCTAACAACATGAC |
| MV19AGO32 SPL6721_g1.2_R | CAAATTTCGTTATGTGGGATTACG |
| MV19AGO33 SPL0991_g3.2&g3.6_F | GTATACATGACACTGTGGCTA |
| MV19AGO34 SPL0991_g3.2&g3.6_R | GTGAAATTTCTCTACTCTGTCAGG |
| MV19AGO35 SPL1538_g3.2&g3.6_F | GAACTGGGTTCAGTTTCTTCTC |
| MV19AGO36 SPL1538_g3.2&g3.6_R | GTTTTGTATCCGCGTATCGACA |
| MV19AGO37 SPL3348_g3.3_F | GAGGAAGACGAAGATGTTGTA |
| MV19AGO38 SPL3348_g3.3_R | CATTTTAAGACAACACAAGCACT |
| MV19AGO39 SPL3942_g3.3_F | GAGGAAGACGAAGATGTTGTG |
| MV19AGO40 SPL3942_g3.3_R | GAATTTAACTTTTGTACACTGACC |
| MV19AGO41 SPL3348_g3.1_F | GCAAGGATGTTGTAACTTGGTTC |
| MV19AGO42 SPL3348_g3.1_R | ACAGACAAAGTTAATGACACACA |
| MV19AGO43 SPL3942_g3.1_F | GCAAGGATGTTGTATCTTGGTTG |
| MV19AGO44 SPL3942_g3.1_R | ACAGACAAAGTTAATGACTCACT |
| JS19ENE01_ALS_F/seq | CAACGTCTTTGCGTACCCAGG |
| JS19ENE02_ALS_R | CCCCAAATGCGAGCAACAAATC |
| JS19ENE03_Nos_F/seq | TTGAAGGAGCCACTGAGCC |

| | |
|---|---|
| JS19ENE04_Nos_R | GGCTGCGAAATGCCCATACTG |
| JS19ENE05_Mtb_F/seq | AGTCGCGGTCGATAGAGAATG |
| JS19ENE06_Mtb_R | ACATCGACTGAAATCCCTGG |
| JS19ENE07_35s_F/seq | CGAGGAGCATCGTGGAAAAAG |
| JS19ENE08_35s_R | CAGGCCATTCATCCCATGATTC |
| JO16DIC16_FT_F | CTAGAAAACCTATGGCTATAAGGG |
| JO16DIC17_FT_R | GTTCTCGAGAGGTATAATATAGGC |
| JO16DIC18_FTseq | CACAAGCACGCATAGAAC |
| JO17AB21_CBP_F | TGTCTAGACTGGTGCATTACTTC |
| JO17AB21_CBP_R/seq | GTTGCCAAAAGGATCACTCAAAT |

#### Table S4. Mutations of the MPO and SPL T1 plants described in this work.

**Plant MPO16A-1-6**

Nitab_0000007g0210.1 (no data for *gMPO1.4*)

<table>
<tr><td>gMPO1.6:</td><td>ACTCGAGGAGGCCACCA|CCGCGGCAAAGTCATTTC</td><td>Mut</td><td>Genotype</td></tr>
<tr><td>Allele 1:</td><td>ACTCGAG----------|---CGGCAAAGTCATTTC</td><td>-13</td><td rowspan="2">HM</td></tr>
<tr><td>Allele 2:</td><td>ACTCGAG----------|---CGGCAAAGTCATTTC</td><td>-13</td></tr>
</table>

Nitab_0002631g0030.1 (no data for *gMPO1.4*)

<table>
<tr><td>gMPO1.6 seq:</td><td>ACTCGAGGAGGCCACCA|CCGCGGCAAAGTCATTTCATC</td><td>Mut</td><td>Genotype</td></tr>
<tr><td>Allele 1:</td><td>ACTCGAGGAGGCCAC-A|CCGCGGCAAAGTCATTTCATC</td><td>-1</td><td rowspan="2">HM</td></tr>
<tr><td>Allele 2:</td><td>ACTCGAGGAGGCCAC-A|CCGCGGCAAAGTCATTTCATC</td><td>-1</td></tr>
</table>

Nitab_0000025g0290.1 (no data for *gMPO2.1*)

<table>
<tr><td>gMPO2.2 seq:</td><td>AGGTATACAAATCATGA|CAAGGGCTCAGACTTGCCAT</td><td>Mut</td><td>Genotype</td></tr>
<tr><td>Allele 1:</td><td>AGGTATACAAATCATGA|TCAAGGGCTCAGACTTGCCAT</td><td>+1</td><td rowspan="2">HM</td></tr>
<tr><td>Allele 2:</td><td>AGGTATACAAATCATGA|TCAAGGGCTCAGACTTGCCAT</td><td>+1</td></tr>
</table>

Nitab_0000983g0060.1 (no data for *gMPO2.4*)

<table>
<tr><td>gMPO2.5 seq:</td><td>AGGTATCCAAATCATGA|CAAGGGCTCAAACCTGCCAT</td><td>Mut</td><td>Genotype</td></tr>
<tr><td>Allele 1:</td><td>AGGTATCCAAATCATGA|ACAAGGGCTCAAACCTGCCAT</td><td>+1</td><td rowspan="2">HM</td></tr>
<tr><td>Allele 2:</td><td>AGGTATCCAAATCATGA|ACAAGGGCTCAAACCTGCCAT</td><td>+1</td></tr>
<tr><td>gMPO2.3 seq:</td><td>GCTCGAGGTGGACATCA|CAGGGGAAAAGTCATCGCAT</td><td>Mut</td><td>Genotype</td></tr>
<tr><td>Allele 1:</td><td>GCTCGAGGTGGACATCA|ACAGGGGAAAAGTCATCGCAT</td><td>+1</td><td rowspan="2">HM</td></tr>
<tr><td>Allele 2:</td><td>GCTCGAGGTGGACATCA|ACAGGGGAAAAGTCATCGCAT</td><td>+1</td></tr>
</table>

**Plant MPO16A-1-10**

Nitab_0000007g0210.1 (no data for *gMPO1.4*)

<table>
<tr><td>gMPO1.6:</td><td>ACTCGAGGAGGCCACCA|CCGCGGCAAAGTCATTTC</td><td>Mut</td><td>Genotype</td></tr>
<tr><td>Allele 1:</td><td>ACTCGAG----------|---CGGCAAAGTCATTTC</td><td>-13</td><td rowspan="2">HM</td></tr>
<tr><td>Allele 2:</td><td>ACTCGAG----------|---CGGCAAAGTCATTTC</td><td>-13</td></tr>
</table>

Nitab_0002631g0030.1 (no data for *gMPO1.4*)

<table>
<tr><td>gMPO1.6 seq:</td><td>ACTCGAGGAGGCCACCA|CCGCGGCAAAGTCATTTCATC</td><td>Mut</td><td>Genotype</td></tr>
<tr><td>Allele 1:</td><td>ACTCGAGGAGGCCAC-A|CCGCGGCAAAGTCATTTCATC</td><td>-1</td><td rowspan="2">BA</td></tr>
<tr><td>Allele 2:</td><td>ACTCGAGGAGGCCACCA|CCCGCGGCAAAGTCATTTCATC</td><td>+1</td></tr>
</table>

Nitab_0000025g0290.1 (no data for *gMPO2.1*)

<table>
<tr><td>gMPO2.2 seq:</td><td>AGGTATACAAATCATGA|CAAGGGCTCAGACTTGCCAT</td><td>Mut</td><td>Genotype</td></tr>
<tr><td>Allele 1:</td><td>AGGTATACAAATCATGA|TCAAGGGCTCAGACTTGCCAT</td><td>+1</td><td rowspan="2">HM</td></tr>
<tr><td>Allele 2:</td><td>AGGTATACAAATCATGA|TCAAGGGCTCAGACTTGCCAT</td><td>+1</td></tr>
</table>

Nitab_0000983g0060.1 (no data for *gMPO2.4*)

<table>
<tr><td>gMPO2.5 seq:</td><td>AGGTATCCAAATCATGA|CAAGGGCTCAAACCTGCCAT</td><td>Mut</td><td>Genotype</td></tr>
<tr><td>Allele 1:</td><td>AGGTATCCAAATCAT--|CAAGGGCTCAAACCTGCCAT</td><td>-2</td><td rowspan="2">BA</td></tr>
<tr><td>Allele 2:</td><td>AGGTATCCAAATCATGA|ACAAGGGCTCAAACCTGCCAT</td><td>+1</td></tr>
<tr><td>gMPO2.3 seq:</td><td>GCTCGAGGTGGACATCA|CAGGGGAAAAGTCATCGCAT</td><td>Mut</td><td>Genotype</td></tr>
<tr><td>Allele 1:</td><td>GCTCGAGGTGGACATCA|ACAGGGGAAAAGTCATCGCAT</td><td>+1</td><td rowspan="2">HM</td></tr>
<tr><td>Allele 2:</td><td>GCTCGAGGTGGACATCA|ACAGGGGAAAAGTCATCGCAT</td><td>+1</td></tr>
</table>

**Plant MPO24-1-4**

Nitab_0000003g0720.1 (no data for *gMPO1.1*)

| *gMPO1.2 seq:* | AGCTGCCGGTGAAACAC\|CCGAGGTTCTTAATCACTA | Mut | Genotype |
|---|---|---|---|
| Allele 1: | AGCTGCCGGTGAAACAC\|-CGAGGTTCTTAATCACTA | -1 | HM |
| Allele 2: | AGCTGCCGGTGAAACAC\|-CGAGGTTCTTAATCACTA | -1 | |
| *gMPO1.3 seq:* | GCTCGAGGTGGACATCA\|CAAGGGAAAAGTGATTTCA | Mut | Genotype |
| Allele 1: | GCT--------------\|CAAGGGAAAAGTGATTTCA | -14 | HM |
| Allele 2: | GCT--------------\|CAAGGGAAAAGTGATTTCA | -14 | |

Nitab_0000007g0210.1 (no data for *gMPO1.4*)

| *gMPO1.6:* | TGAGACCAGCATATGGATTGTTGAGCTCTCGGAGGTTC ATGCTGTGACTCGAGGAGGCCACCA\|CCGCGGCAAAG | Mut | Genotype |
|---|---|---|---|
| Allele 1: | TGAGACCAGCATATGGATTGTTGAGCTCTCGGAGGTTC ATGCTGTGACTCGAGGAGGCCACC-\|CCGCGGCAAAG | -1 | BA |
| Allele 2: | TGAGACC------------------------------- -------------------------\|TCCGCGGCAAAG | -56/+1 | |

Nitab_0002631g0030.1 (no data for *gMPO1.4*)

| *gMPO1.6 seq:* | ACTCGAGGAGGCCACCA\|CCGCGGCAAAGTCATTTCATC | Mut | Genotype |
|---|---|---|---|
| Allele 1: | ACTCGAGGAGGCCACCA\|TCCGCGGCAAAGTCATTTCATC | +1 | HM |
| Allele 2: | ACTCGAGGAGGCCACCA\|TCCGCGGCAAAGTCATTTCATC | +1 | |

Nitab_0000025g0290.1 (no data for *gMPO2.1*)

| *gMPO2.2 seq:* | AGGTATACAAATCATGA\|CAAGGGCTCAGACTTGCCAT | Mut | Genotype |
|---|---|---|---|
| Allele 1: | AGGTATACAAATCATGA\|ACAAGGGCTCAGACTTGCCAT | +1 | HM |
| Allele 2: | AGGTATACAAATCATGA\|ACAAGGGCTCAGACTTGCCAT | +1 | |
| *gMPO2.3 seq:* | GCTCGAGGTGGACATCA\|CAGGGGAAAAGTCATCTCAT | Mut | Genotype |
| Allele 1: | GCTCGAGGTGGACATCA\|ACAGGGGAAAAGTCATCTCAT | +1 | HM |
| Allele 2: | GCTCGAGGTGGACATCA\|ACAGGGGAAAAGTCATCTCAT | +1 | |

Nitab_0000983g0060.1 (no data for *gMPO2.4*)

| *gMPO2.5 seq:* | AGGTATCCAAATCATGA\|CAAGGGCTCAAACCTGCCAT | Mut | Genotype |
|---|---|---|---|
| Allele 1: | AGGTATCCAAATCATG-\|-AAGGGCTCAAACCTGCCAT | -2 | HM |
| Allele 2: | AGGTATCCAAATCATG-\|-AAGGGCTCAAACCTGCCAT | -2 | |

**Plant MPO24-1-6**

Nitab_0000003g0720.1 (no data for *gMPO1.1*)

| *gMPO1.2 seq:* | AGCTGCCGGTGAAACAC\|CCGAGGTTCTTAATCACTA | Mut | Genotype |
|---|---|---|---|
| Allele 1: | AGCTGCCGGTGAAACAC\|CCGAGGTTCTTAATCACTA | WT | HT |
| Allele 2: | AGCTGCCGGTGAAACAC\|-CGAGGTTCTTAATCACTA | -1 | |
| *gMPO1.3 seq:* | GCTCGAGGTGGACATCA\|CAAGGGAAAAGTGATTTCA | Mut | Genotype |
| Allele 1: | GCTCGAGGTGGACATCA\|ACAAGGGAAAAGTGATTTCA | +1 | BA |
| Allele 2: | GCT--------------\|CAAGGGAAAAGTGATTTCA | -14 | |

Nitab_0002358g0050.1 (no data for *gMPO1.1*)

| *gMPO1.3 seq:* | GCTCGAGGTGGACATCA\|CAAGGGAAAAGTGATTTCA | Mut | Genotype |
|---|---|---|---|
| Allele 1: | GCTCGAGGTGGACATCA\|CAAGGGAAAAGTGATTTCA | WT | HT |
| Allele 2: | GCTCGAGGTGGACATCA\|ACAAGGGAAAAGTGATTTCA | +1 | |

Nitab_0000007g0210.1 (no data for *gMPO1.4*)

| *gMPO1.6:* | TGAGACCAGCATATGGATTGTTGAGCTCTCGGAGGTTC ATGCTGTGACTCGAGGAGGCCACCA\|CCGCGGCAAAG | Mut | Genotype |
|---|---|---|---|

| | | | |
|---|---|---|---|
| Allele 1: | TGAGACCAGCATATGGATTGTTGAGCTCTCGGAGGTTC<br>ATGCTGTGACTCGAGGAGGCCACC-\|CCGCGGCAAAG | -1 | BA |
| Allele 2: | TGAGACC-------------------------------<br>-------------------------\|TCCGCGGCAAAG | -56/+1 | |

Nitab_0002631g0030.1 (no data for *gMPO1.4*)

| *gMPO1.6 seq:* | ACTCGAGGAGGCCACCA\|CCGCGGCAAAGTCATTTCATC | Mut | Genotype |
|---|---|---|---|
| Allele 1: | ACTCGAGGAGGCCACCA\|TCCGCGGCAAAGTCATTTCATC | +1 | BA |
| Allele 2: | ACTCGAGGAGGC-----\|CCGCGGCAAAGTCATTTCATC | -5 | |

Nitab_0000025g0290.1 (no data for *gMPO2.1*)

| *gMPO2.2 seq:* | AGGTATACAAATCATGA\|CAAGGGCTCAGACTTGCCAT | Mut | Genotype |
|---|---|---|---|
| Allele 1: | AGGTATACAAATCATGA\|ACAAGGGCTCAGACTTGCCAT | +1 | HM |
| Allele 2: | AGGTATACAAATCATGA\|ACAAGGGCTCAGACTTGCCAT | +1 | |
| *gMPO2.3 seq:* | GCTCGAGGTGGACATCA\|CAGGGGAAAAGTCATCTCAT | Mut | Genotype |
| Allele 1: | GCTCGAGGTGGACATCA\|ACAGGGGAAAAGTCATCTCAT | +1 | HM |
| Allele 2: | GCTCGAGGTGGACATCA\|ACAGGGGAAAAGTCATCTCAT | +1 | |

Nitab_0000983g0060.1 (no data for gMPO2.3)

| *gMPO2.5 seq:* | AGGTATCCAAATCATGA\|CAAGGGCTCAAACCTGCCAT | Mut | Genotype |
|---|---|---|---|
| Allele 1: | AGGTATCCAAATCATG-\|-AAGGGCTCAAACCTGCCAT | -2 | HM |
| Allele 2: | AGGTATCCAAATCATG-\|-AAGGGCTCAAACCTGCCAT | -2 | |

**Plant MPO24-1-7**

Nitab_0000003g0720.1 (no data for *gMPO1.1*)

| *gMPO1.3 seq:* | GCTCGAGGTGGACATCA\|CAAGGGAAAAGTGATTTCA | Mut | Genotype |
|---|---|---|---|
| Allele 1: | GCTCGAGGTGGACATCA\|ACAAGGGAAAAGTGATTTCA | +1 | HM |
| Allele 2: | GCTCGAGGTGGACATCA\|ACAAGGGAAAAGTGATTTCA | +1 | |

Nitab_0002358g0050.1 (no data for *gMPO1.1*)

| *gMPO1.3 seq:* | GCTCGAGGTGGACATCA\|CAAGGGAAAAGTGATTTC | Mut | Genotype |
|---|---|---|---|
| Allele 1: | GCTCGAGGTGGACATCA\|ACAAGGGAAAAGTGATTTC | +1 | HM |
| Allele 2: | GCTCGAGGTGGACATCA\|ACAAGGGAAAAGTGATTTC | +1 | |

Nitab_0000007g0210.1 (no data for *gMPO1.4*)

| *gMPO1.6:* | TGAGACCAGCATATGGATTGTTGAGCTCTCGGAGGTTC<br>ATGCTGTGACTCGAGGAGGCCACCA\|CCGCGGCAAAG | Mut | Genotype |
|---|---|---|---|
| Allele 1: | TGAGACC-------------------------------<br>-------------------------\|TCCGCGGCAAAG | -56/+1 | HM |
| Allele 2: | TGAGACC-------------------------------<br>-------------------------\|TCCGCGGCAAAG | -56/+1 | |

Nitab_0002631g0030.1 (no data for *gMPO1.4*)

| *gMPO1.6 seq:* | ACTCGAGGAGGCCACCA\|CCGCGGCAAAGTCATTTCATC | Mut | Genotype |
|---|---|---|---|
| Allele 1: | ACTCGAGGAGGCCACCA\|TCCGCGGCAAAGTCATTTCATC | +1 | HM |
| Allele 2: | ACTCGAGGAGGCCACCA\|TCCGCGGCAAAGTCATTTCATC | +1 | |

Nitab_0000025g0290.1 (no data for *gMPO2.1*)

| *gMPO2.2 seq:* | AGGTATACAAATCATGA\|CAAGGGCTCAGACTTGCCAT | Mut | Genotype |
|---|---|---|---|
| Allele 1: | AGGTATACAAATCATGA\|ACAAGGGCTCAGACTTGCCAT | +1 | HM |
| Allele 2: | AGGTATACAAATCATGA\|ACAAGGGCTCAGACTTGCCAT | +1 | |

| *gMPO2.3 seq:* | GCTCGAGGTGGACATCA\|CAGGGGAAAAGTCATCTCAT | Mut | Genotype |
|---|---|---|---|
| Allele 1: | GCTCGAGGTGGACATCA\|ACAGGGGAAAAGTCATCTCAT | +1 | HM |
| Allele 2: | GCTCGAGGTGGACATCA\|ACAGGGGAAAAGTCATCTCAT | +1 | |

Nitab_0000983g0060.1 (no data for *gMPO2.4*)

| *gMPO2.5 seq:* | AGGTATCCAAATCATGA\|CAAGGGCTCAAACCTGCCAT | Mut | Genotype |
|---|---|---|---|
| Allele 1: | AGGTATCCAAATCATG-\|-AAGGGCTCAAACCTGCCAT | -2 | HM |
| Allele 2: | AGGTATCCAAATCATG-\|-AAGGGCTCAAACCTGCCAT | -2 | |

**Plant MPO25-2-2**

Nitab_0000003g0720.1 (no data for *gMPO1.1*)

| *gMPO1.3 seq:* | GCTCGAGGTGGACATCA\|CAAGGGAAAAGTGATTTCA | Mut | Genotype |
|---|---|---|---|
| Allele 1: | GCTCGAGGTGGACATCA\|ACAAGGGAAAAGTGATTTCA | +1 | BA |
| Allele 2: | GCT--------------\|CAAGGGAAAAGTGATTTCA | -14 | |

Nitab_0000007g0210.1 (no data for *gMPO1.4*)

| *gMPO1.6:* | ACTCGAGGAGGCCACCA\|CCGCGGCAAAG | Mut | Genotype |
|---|---|---|---|
| Allele 1: | ACTCGAGGAGGCCACC-\|CCGCGGCAAAG | -1 | HM |
| Allele 2: | ACTCGAGGAGGCCACC-\|CCGCGGCAAAG | -1 | |

Nitab_0002631g0030.1 (no data for *gMPO1.4*)

| *gMPO1.6 seq:* | ACTCGAGGAGGCCACCA\|CCGCGGCAAAGTCATTTCATC | Mut | Genotype |
|---|---|---|---|
| Allele 1: | ACTCGAGGAGGC-----\|CCGCGGCAAAGTCATTTCATC | -5 | HM |
| Allele 2: | ACTCGAGGAGGC-----\|CCGCGGCAAAGTCATTTCATC | -5 | |

Nitab_0000025g0290.1 (no data for *gMPO2.1)*

| *gMPO2.2 seq:* | AGGTATACAAATCATGA\|CAAGGGCTCAGACTTGCCAT | Mut | Genotype |
|---|---|---|---|
| Allele 1: | AGGTATACAAATCATGA\|ACAAGGGCTCAGACTTGCCAT | +1 | HM |
| Allele 2: | AGGTATACAAATCATGA\|ACAAGGGCTCAGACTTGCCAT | +1 | |
| *gMPO2.3 seq:* | GCTCGAGGTGGACATCA\|CAGGGGAAAAGTCATCTCAT | Mut | Genotype |
| Allele 1: | GCTCGAGGTGGACATCA\|ACAGGGGAAAAGTCATCTCAT | +1 | HM |
| Allele 2: | GCTCGAGGTGGACATCA\|ACAGGGGAAAAGTCATCTCAT | +1 | |

Nitab_0000983g0060.1 (no data for *gMPO2.4*)

| *gMPO2.3 seq:* | GCTCGAGGTGGACATCA\|CAGGGGAAAAGTCATCGCATC CAATGTTGT | Mut | Genotype |
|---|---|---|---|
| Allele 1: | GCTCGAGGTGGATTGATGTCCAATGTTGTCCCCTGATGT CCAAATGCTTGTCTCCAAATACTATGGGTGCGTGCATCC AATGTTGT | -20/+60 | HM |
| Allele 2: | GCTCGAGGTGGATTGATGTCCAATGTTGTCCCCTGATGT CCAAATGCTTGTCTCCAAATACTATGGGTGCGTGCATCC AATGTTGT | -20/+60 | |
| *gMPO2.5 seq:* | AGGTATCCAAATCATGA\|CAAGGGCTCAAACCTGCCAT | Mut | Genotype |
| Allele 1: | AGGTATCCAAATCATGA\|TCAAGGGCTCAAACCTGCCAT | +1 | HM |
| Allele 2: | AGGTATCCAAATCATGA\|TCAAGGGCTCAAACCTGCCAT | +1 | |

**Plant SPL2-1**

Nitab_0002299g0030.1

<table>
<tr><td>gSPL1.6 seq:</td><td>ATGGACATAACAGGCGT|CGAAGGAAACCTCAGCC</td><td>Mut</td><td>Genotype</td></tr>
<tr><td>Allele 1:</td><td>ATGGACATAACAGGCGT|ACGAAGGAAACCTCAGCC</td><td>+1</td><td rowspan="2">BA</td></tr>
<tr><td>Allele 2:</td><td>ATGGA------------|-----------TCAGCC</td><td>-23</td></tr>
</table>

Nitab_0001010g0010.1

<table>
<tr><td>gSPL1.6 seq:</td><td>ATGGACATAACAGGCGT|CGAAGGAAACCTCAGCC</td><td>Mut</td><td>Genotype</td></tr>
<tr><td>Allele 1:</td><td>ATGGACATAACAGGCGT|TCGAAGGAAACCTCAGCC</td><td>+1</td><td rowspan="2">HM</td></tr>
<tr><td>Allele 2:</td><td>ATGGACATAACAGGCGT|TCGAAGGAAACCTCAGCC</td><td>+1</td></tr>
</table>

Nitab_0000861g0050.1

<table>
<tr><td>gSPL2.4 seq:</td><td>TGTGATCAGAGAGCCTC|CTGCGGCAGCTTCTCT</td><td>Mut</td><td>Genotype</td></tr>
<tr><td>Allele 1:</td><td>TGTGATCAGAGAGCCT-|CTGCGGCAGCTTCTCT</td><td>-1</td><td rowspan="2">HM</td></tr>
<tr><td>Allele 2:</td><td>TGTGATCAGAGAGCCT-|CTGCGGCAGCTTCTCT</td><td>-1</td></tr>
<tr><td>gSPL3.5 seq:</td><td>AGATGGAGTGGAATGCA|AAGTGGGACTGGGGAAA</td><td>Mut</td><td>Genotype</td></tr>
<tr><td>Allele 1:</td><td>AGATGGAGTGGAATGCA|AAGTGGGACTGGGGAAA</td><td>WT</td><td rowspan="2">HT</td></tr>
<tr><td>Allele 2:</td><td>AGATGGAGTGGAATGCA|AAAGTGGGACTGGGGAAA</td><td>+1</td></tr>
</table>

**Plant SPL2-3**

Nitab_0003942g0050.1

<table>
<tr><td>gSPL3.1 seq:</td><td>TTAAAGGGGCATCAGTC|CAATGGCAGAGAAATC</td><td>Mut</td><td>Genotype</td></tr>
<tr><td>Allele 1:</td><td>TTAAAGGGGCATCAGTC|CCAATGGCAGAGAAATC</td><td>+1</td><td rowspan="2">HM</td></tr>
<tr><td>Allele 2:</td><td>TTAAAGGGGCATCAGTC|CCAATGGCAGAGAAATC</td><td>+1</td></tr>
</table>

Nitab_0001538g0080.1

<table>
<tr><td>gSPL3.6 seq:</td><td>CCTTCAACTTGACACCT|AGGTGGCTGACCACCCT</td><td>Mut</td><td>Genotype</td></tr>
<tr><td>Allele 1:</td><td>CCTTCAACTTGA----T|AGGTGGCTGACCACCCT</td><td>-4</td><td rowspan="2">HM</td></tr>
<tr><td>Allele 2:</td><td>CCTTCAACTTGA----T|AGGTGGCTGACCACCCT</td><td>-4</td></tr>
</table>

Nitab_0002299g0030.1

<table>
<tr><td>gSPL1.6 seq:</td><td>ATGGACATAACAGGCGT|CGAAGGAAACCTCAGCC</td><td>Mut</td><td>Genotype</td></tr>
<tr><td>Allele 1:</td><td>ATGGACATAACAGGCGT|ACGAAGGAAACCTCAGCC</td><td>+1</td><td rowspan="2">HM</td></tr>
<tr><td>Allele 2:</td><td>ATGGACATAACAGGCGT|ACGAAGGAAACCTCAGCC</td><td>+1</td></tr>
</table>

Nitab_0001010g0010.1

<table>
<tr><td>gSPL1.6 seq:</td><td>ATGGACATAACAGGCGT|CGAAGGAAACCTCAGCC</td><td>Mut</td><td>Genotype</td></tr>
<tr><td>Allele 1:</td><td>ATGGACATAACAGGCGT|TCGAAGGAAACCTCAGCC</td><td>+1</td><td rowspan="2">HM</td></tr>
<tr><td>Allele 2:</td><td>ATGGACATAACAGGCGT|TCGAAGGAAACCTCAGCC</td><td>+1</td></tr>
</table>

Nitab_0000067g0130.1

<table>
<tr><td>gSPL2.1 seq:</td><td>CGAACAGTTGAAAGCAA|CAGTGGTGGAAGCAGT</td><td>Mut</td><td>Genotype</td></tr>
<tr><td>Allele 1:</td><td>CGAACAGTTGAAAGCAA|ACAGTGGTGGAAGCAGT</td><td>+1</td><td rowspan="2">HM</td></tr>
<tr><td>Allele 2:</td><td>CGAACAGTTGAAAGCAA|ACAGTGGTGGAAGCAGT</td><td>+1</td></tr>
</table>

Nitab_0000861g0050.1

<table>
<tr><td>gSPL3.5 seq:</td><td>AGATGGAGTGGAATGCA|AAGTGGGACTGGGGAAA</td><td>Mut</td><td>Genotype</td></tr>
<tr><td>Allele 1:</td><td>AGATGGAGTGGAATGCA|AAAGTGGGACTGGGGAAA</td><td>+1</td><td rowspan="2">HM</td></tr>
<tr><td>Allele 2:</td><td>AGATGGAGTGGAATGCA|AAAGTGGGACTGGGGAAA</td><td>+1</td></tr>
</table>

**Plant SPL2-5**

Nitab_0003942g0050.1

| *gSPL3.1 seq:* | TTAAAGGGGCATCAGTC\|CAATGGCAGAGAAATC | Mut | Genotype |
|---|---|---|---|
| Allele 1: | TTAAAGGGGCATCAGTC\|CAATGGCAGAGAAATC | WT | HT |
| Allele 2: | TTAAAGGGGCATCAGTC\|CCAATGGCAGAGAAATC | +1 | |

Nitab_0001538g0080.1

| *gSPL3.6 seq:* | CCTTCAACTTGACACCT\|AGGTGGCTGACCACCCT | Mut | Genotype |
|---|---|---|---|
| Allele 1: | CCTTCAACTTGA----T\|AGGTGGCTGACCACCCT | -4 | HM |
| Allele 2: | CCTTCAACTTGA----T\|AGGTGGCTGACCACCCT | -4 | |

Nitab_0002299g0030.1

| *gSPL1.6 seq:* | ATGGACATAACAGGCGT\|CGAAGGAAACCTCAGCC | Mut | Genotype |
|---|---|---|---|
| Allele 1: | ATGGACATAACAGGCGT\|ACGAAGGAAACCTCAGCC | +1 | HM |
| Allele 2: | ATGGACATAACAGGCGT\|ACGAAGGAAACCTCAGCC | +1 | |

Nitab_0001010g0010.1

| *gSPL1.6 seq:* | ATGGACATAACAGGCGT\|CGAAGGAAACCTCAGCC | Mut | Genotype |
|---|---|---|---|
| Allele 1: | ATGGACATAACAGGCGT\|TCGAAGGAAACCTCAGCC | +1 | HM |
| Allele 2: | ATGGACATAACAGGCGT\|TCGAAGGAAACCTCAGCC | +1 | |

Nitab_0000067g0130.1

| *gSPL2.1 seq:* | CGAACAGTTGAAAGCAA\|CAGTGGTGGAAGCAGT | Mut | Genotype |
|---|---|---|---|
| Allele 1: | CGAACAGTTGAAAGCAA\|CAGTGGTGGAAGCAGT | WT | HT |
| Allele 2: | CGAACAGTTGAAAGCAA\|ACAGTGGTGGAAGCAGT | +1 | |

Nitab_0000861g0050.1

| *gSPL2.4 seq:* | TGTGATCAGAGAGCCTC\|CTGCGGCAGCTTCTCTTCTTTT | Mut | Genotype |
|---|---|---|---|
| Allele 1: | TGTGATCAGAGAGCCT-\|CTGCGGCAGCTTCTCTTCTTTT | -1 | HM |
| Allele 2: | TGTGATCAGAGAGCCT-\|CTGCGGCAGCTTCTCTTCTTTT | -1 | |
| *gSPL3.5 seq:* | AGATGGAGTGGAATGCA\|AAGTGGGACTGGGGAAA | Mut | Genotype |
| Allele 1: | AGATGGAGTGGAATGCA\|AAGTGGGACTGGGGAAA | WT | HT |
| Allele 2: | AGATGGAGTGGAATGCA\|AAAGTGGGACTGGGGAAA | +1 | |

**Plant SPL10-3**

Nitab_0003942g0050.1

| *gSPL3.1 seq:* | TTAAAGGGGCATCAGTC\|CAATGGCAGAGAAATC | Mut | Genotype |
|---|---|---|---|
| Allele 1: | TTAAAGGGGCATCAGTC\|CAATGGCAGAGAAATC | WT | HT |
| Allele 2: | TTAAAGGGGCATCAGTC\|TCAATGGCAGAGAAATC | +1 | |

Nitab_0003348g0050.1

| *gSPL3.1 seq:* | TTAAAGGGGCATCAGTC\|CAATGGCAGAGAAATC | Mut | Genotype |
|---|---|---|---|
| Allele 1: | TTAAAGGGGCATCAGTC\|TCAATGGCAGAGAAATC | +1 | HM |
| Allele 2: | TTAAAGGGGCATCAGTC\|TCAATGGCAGAGAAATC | +1 | |

Nitab_0001538g0080.1

| *gSPL3.6 seq:* | CCTTCAACTTGACACCT\|AGGTGGCTGACCACCCT | Mut | Genotype |
|---|---|---|---|
| Allele 1: | CCTTCAACTTGACAC-A\|AGGTGGCTGACCACCCT | -2/+1 | BA |
| Allele 2: | CCTTCAACTTGACAC--\|AGGTGGCTGACCACCCT | -2 | |
| *gSPL3.2 seq:* | ATCTACTTTGAAAATG\|TGGTGGGTCATCGCCGGT | Mut | Genotype |
| Allele 1: | ATCTACTTTG------\|-GGTGGGTCATCGCCGGT | -7 | BA |
| Allele 2: | ATCTACTTTGAAAATG\|TAGGTGGGTCATCGCCGGT | +1 | |

Nitab_0000991g0020.1

| *gSPL3.6 seq:* | CCTTCAACTTGACACCT\|AGGTGGCTGACCACCCTGT | Mut | Genotype |
|---|---|---|---|
| Allele 1: | CCTTCAACTTGACACCT\|AAGGTGGCTGACCACCCTGT | +1 | HM |
| Allele 2: | CCTTCAACTTGACACCT\|AAGGTGGCTGACCACCCTGT | +1 | |

Nitab_0002299g0030.1

| *gSPL1.6 seq:* | ATGGACATAACAGGCGT\|CGAAGGAAACCTCAGCC | Mut | Genotype |
|---|---|---|---|
| Allele 1: | ATGGACATAACAGGCGT\|TCGAAGGAAACCTCAGCC | +1 | BA |
| Allele 2: | ATGGACATAACAGGCGT\|ACGAAGGAAACCTCAGCC | +1 | |

Nitab_0001010g0010.1

| *gSPL1.6 seq:* | ATGGACATAACAGGCGT\|CGAAGGAAACCTCAGCC | Mut | Genotype |
|---|---|---|---|
| Allele 1: | ATGGACATAACAGGCGT\|TCGAAGGAAACCTCAGCC | +1 | HM |
| Allele 2: | ATGGACATAACAGGCGT\|TCGAAGGAAACCTCAGCC | +1 | |

Nitab_0001315g0110.1

| *gSPL3.5 seq:* | AGATGGAGTGGAATGCA\|AAGTGGGACTGGGGAAACCA | Mut | Genotype |
|---|---|---|---|
| Allele 1: | AGATGGAGTGGAATGCA\|AAAGTGGGACTGGGGAAACCA | +1 | HM |
| Allele 2: | AGATGGAGTGGAATGCA\|AAAGTGGGACTGGGGAAACCA | +1 | |

Nitab_0003900g0020.1

| *gSPL2.5 seq:* | GGAAACCATCCAGTTCA\|ACTCGGCAAGGCTTTCTTCAT | Mut | Genotype |
|---|---|---|---|
| Allele 1: | GGAAACCATCCAGTT--\|ACTCGGCAAGGCTTTCTTCAT | -2 | BA |
| Allele 2: | GGAAACCATCCAG---A\|ACTCGGCAAGGCTTTCTTCAT | -3 | |

Nitab_0000861g0050.1

| *gSPL2.4 seq:* | TGTGATCAGAGAGCCTC\|CTGCGGCAGCTTCTCTTCTTT | Mut | Genotype |
|---|---|---|---|
| Allele 1: | TGTGATCAGAGAG---C\|CTGCGGCAGCTTCTCTTCTTT | -3 | HM |
| Allele 2: | TGTGATCAGAGAG---C\|CTGCGGCAGCTTCTCTTCTTT | -3 | |

Nitab_0007217g0040.1

| *gSPL2.3 seq:* | TAGAAATTCACCCCCCA\|TGGAGGCTTTGGTAGGCT | Mut | Genotype |
|---|---|---|---|
| Allele 1: | TAGAAATT---------\|TGGAGGCTTTGGTAGGCT | -9 | HM |
| Allele 2: | TAGAAATT---------\|TGGAGGCTTTGGTAGGCT | -9 | |

**Plant SPL10-5**

Nitab_0003942g0050.1

| *gSPL3.1 seq:* | TTAAAGGGGCATCAGTC\|CAATGGCAGAGAAATC | Mut | Genotype |
|---|---|---|---|
| Allele 1: | TTAAAGGGGCATCAGTC\|TCAATGGCAGAGAAATC | +1 | HM |
| Allele 2: | TTAAAGGGGCATCAGTC\|TCAATGGCAGAGAAATC | +1 | |
| gSPL3.3 seq: | GAAATTCACAGACCTTG\|TGGCGGCGATGGTATGTC | Mut | Genotype |
| Allele 1: | GAAATTCACAGACCTT-\|TGGCGGCGATGGTATGTC | -1 | HM |
| Allele 2: | GAAATTCACAGACCTT-\|TGGCGGCGATGGTATGTC | -1 | |

Nitab_0001538g0080.1

| *gSPL3.6 seq:* | CCTTCAACTTGACACCT\|AGGTGGCTGACCACCCT | Mut | Genotype |
|---|---|---|---|
| Allele 1: | CCTTCAACTTGACAC-A\|AGGTGGCTGACCACCCT | -2/+1 | HM |
| Allele 2: | CCTTCAACTTGACAC-A\|AGGTGGCTGACCACCCT | -2/+1 | |

Nitab_0000991g0020.1

| *gSPL3.6 seq:* | CCTTCAACTTGACACCT\|AGGTGGCTGACCACCCTGT | Mut | Genotype |
|---|---|---|---|
| Allele 1: | CCTTCAACTTGACACCT\|AGGTGGCTGACCACCCTGT | WT | HT |

| | | |
|---|---|---|
| Allele 2: | CCTTCAACTTGACACCT\|TAGGTGGCTGACCACCCTGT | +1 |

Nitab_0002299g0030.1

| *gSPL1.6 seq:* | ATGGACATAACAGGCGT\|CGAAGGAAACCTCAGCC | Mut | Genotype |
|---|---|---|---|
| Allele 1: | ATGGACATAACAGGCGT\|TCGAAGGAAACCTCAGCC | +1 | HM |
| Allele 2: | ATGGACATAACAGGCGT\|TCGAAGGAAACCTCAGCC | +1 | |

Nitab_0001010g0010.1

| *gSPL1.6 seq:* | ATGGACATAACAGGCGT\|CGAAGGAAACCTCAGCC | Mut | Genotype |
|---|---|---|---|
| Allele 1: | ATGGACATAACAGGCGT\|TCGAAGGAAACCTCAGCC | +1 | BA |
| Allele 2: | ATGGACATAACAGGCGT\|CCGAAGGAAACCTCAGCC | +1 | |

Nitab_0003900g0020.1

| *gSPL2.5 seq:* | GGAAACCATCCAGTTCA\|ACTCGGCAAGGCTTTCTTCAT | Mut | Genotype |
|---|---|---|---|
| Allele 1: | GGAAACCATCCAG---A\|ACTCGGCAAGGCTTTCTTCAT | -3 | HM |
| Allele 2: | GGAAACCATCCAG---A\|ACTCGGCAAGGCTTTCTTCAT | -3 | |

Nitab_0000861g0050.1

| *gSPL2.4rc & 2.5 seq:* | CCGCAGGAGGCTCTCTGATCACAATGCACGACGCCGCAA ACCACAGCAGGAAACCATCCAGTTCAACTCGGCAAGGCT | Mut | Genotype |
|---|---|---|---|
| Allele 1: | CCGCAG----------------------------------- ----------------------------ACTCGGCAAGGCT | -59 | HM |
| Allele 2: | CCGCAG----------------------------------- ----------------------------ACTCGGCAAGGCT | -59 |  |

Nitab_0007217g0040.1

| *gSPL2.3 seq:* | TAGAAATTCACCCCCCA\|TGGAGGCTTTGGTAGGCT | Mut | Genotype |
|---|---|---|---|
| Allele 1: | TAGAAATTCACCCCCCA\|TGGAGGCTTTGGTAGGCT | WT | HT |
| Allele 2: | TAGAAATT---------\|TGGAGGCTTTGGTAGGCT | -9 | |

**Plant SPL11-3**

Nitab_0004959g0040.1

| *gSPL1.3 seq:* | AAGGGTCAAGTACTTCA\|ATGAGGTGTTGCCAAGCTGA | Mut | Genotype |
|---|---|---|---|
| Allele 1: | AAGGGTCAAGTACTTCA\|ATGAGGTGTTGCCAAGCTGA | WT | HT |
| Allele 2: | AAGGGTCAAGTACTTCA\|AATGAGGTGTTGCCAAGCTGA | +1 | |

Nitab_0002299g0030.1

| *gSPL1.6 seq:* | ATGGACATAACAGGCGT\|CGAAGGAAACCTCAGCC | Mut | Genotype |
|---|---|---|---|
| Allele 1: | ATGGACATAACAGGCGT\|TCGAAGGAAACCTCAGCC | +1 | BA |
| Allele 2: | ATGGACATAACAGGCGT\|ACGAAGGAAACCTCAGCC | +1 | |

Nitab_0001010g0010.1

| *gSPL1.6 seq:* | ATGGACATAACAGGCGT\|CGAAGGAAACCTCAGCC | Mut | Genotype |
|---|---|---|---|
| Allele 1: | ATGGACATAACAGGCGT\|ACGAAGGAAACCTCAGCC | +1 | HM |
| Allele 2: | ATGGACATAACAGGCGT\|ACGAAGGAAACCTCAGCC | +1 | |

Nitab_0000861g0050.1

| *gSPL2.4 seq:* | TGTGATCAGAGAGCCTC\|CTGCGGCAGCTTCTCTTCTTT | Mut | Genotype |
|---|---|---|---|
| Allele 1: | TGTGATCAGAGAGCCTC\|ACTGCGGCAGCTTCTCTTCTTT | +1 | HM |
| Allele 2: | TGTGATCAGAGAGCCTC\|ACTGCGGCAGCTTCTCTTCTTT | +1 | |

**Plant SPL11-5**

Nitab_0003942g0050.1

| *gSPL3.3 seq:* | GAAATTCACAGACCTTG\|TGGCGGCGATGGTATGTC | Mut | Genotype |
|---|---|---|---|
| Allele 1: | GAAATTCACAGACCTTG\|TGGCGGCGATGGTATGTC | WT | HT |
| Allele 2: | GAAATTCACAGACCTTG\|TTGGCGGCGATGGTATGTC | +1 | |

Nitab_0002299g0030.1

| *gSPL1.6 seq:* | ATGGACATAACAGGCGT\|CGAAGGAAACCTCAGCC | Mut | Genotype |
|---|---|---|---|
| Allele 1: | ATGGACATAACAGGCGT\|TCGAAGGAAACCTCAGCC | +1 | BA |
| Allele 2: | ATGGACATAACAGGCGT\|ACGAAGGAAACCTCAGCC | +1 | |

Nitab_0001010g0010.1

| *gSPL1.6 seq:* | ATGGACATAACAGGCGT\|CGAAGGAAACCTCAGCC | Mut | Genotype |
|---|---|---|---|
| Allele 1: | ATGGACATAACAGGCGT\|GCGAAGGAAACCTCAGCC | +1 | HM |
| Allele 2: | ATGGACATAACAGGCGT\|GCGAAGGAAACCTCAGCC | +1 | |

Nitab_0003900g0020.1

| *gSPL2.5 seq:* | GGAAACCATCCAGTTCA\|ACTCGGCAAGGCTTTCTTCAT | Mut | Genotype |
|---|---|---|---|
| Allele 1: | GGAAACCATCCAGTTCA\|ACTCGGCAAGGCTTTCTTCAT | WT | HT |
| Allele 2: | GGAAACCATCCAGTTCA\|AACTCGGCAAGGCTTTCTTCAT | +1 | |

**Plant SPL11-6**

Nitab_0003942g0050.1

| *gSPL3.3 seq:* | GAAATTCACAGACCTTG\|TGGCGGCGATGGTATGTC | Mut | Genotype |
|---|---|---|---|
| Allele 1: | GAAATTCACAGACCTTG\|ATGGCGGCGATGGTATGTC | +1 | HM |
| Allele 2: | GAAATTCACAGACCTTG\|ATGGCGGCGATGGTATGTC | +1 | |

Nitab_0004959g0040.1

| *gSPL1.3 seq:* | AAGGGTCAAGTACTTCA\|ATGAGGTGTTGCCAAGCTGA | Mut | Genotype |
|---|---|---|---|
| Allele 1: | AAGGGTCAAGTACTTCA\|ATGAGGTGTTGCCAAGCTGA | WT | HT |
| Allele 2: | AAGGGTCAAGTACTTCA\|AATGAGGTGTTGCCAAGCTGA | +1 | |

Nitab_0002299g0030.1

| *gSPL1.6 seq:* | ATGGACATAACAGGCGT\|CGAAGGAAACCTCAGCC | Mut | Genotype |
|---|---|---|---|
| Allele 1: | ATGGACATAACAGGCGT\|TCGAAGGAAACCTCAGCC | +1 | HM |
| Allele 2: | ATGGACATAACAGGCGT\|TCGAAGGAAACCTCAGCC | +1 | |

Nitab_0001010g0010.1

| *gSPL1.6 seq:* | ATGGACATAACAGGCGT\|CGAAGGAAACCTCAGCC | Mut | Genotype |
|---|---|---|---|
| Allele 1: | ATGGACATAACAGGCGT\|ACGAAGGAAACCTCAGCC | +1 | HM |
| Allele 2: | ATGGACATAACAGGCGT\|ACGAAGGAAACCTCAGCC | +1 | |

Nitab_0003900g0020.1

| *gSPL2.5 seq:* | GGAAACCATCCAGTTCA\|ACTCGGCAAGGCTTTCTTCAT | Mut | Genotype |
|---|---|---|---|
| Allele 1: | GGAAACCATCCAGTTCA\|AACTCGGCAAGGCTTTCTTCAT | +1 | HM |
| Allele 2: | GGAAACCATCCAGTTCA\|AACTCGGCAAGGCTTTCTTCAT | +1 | |

Nitab_0000861g0050.1

| *gSPL2.4 seq:* | TGTGATCAGAGAGCCTC\|CTGCGGCAGCTTCTCTTCTTT | Mut | Genotype |
|---|---|---|---|

| | | | |
|---|---|---|---|
| Allele 1: | TGTGATCAGAGAGCCTC\|ACTGCGGCAGCTTCTCTTCTTT | +1 | HM |
| Allele 2: | TGTGATCAGAGAGCCTC\|ACTGCGGCAGCTTCTCTTCTTT | +1 | |

**Plant SPL15-1**

Nitab_0003942g0050.1

| *gSPL3.3 seq:* | GAAATTCACAGACCTTG\|TGGCGGCGATGGTATGTC | Mut | Genotype |
|---|---|---|---|
| Allele 1: | GAAATTCACAGACCT--\|TGGCGGCGATGGTATGTC | -2 | HM |
| Allele 2: | GAAATTCACAGACCT--\|TGGCGGCGATGGTATGTC | -2 | |

Nitab_0003348g0050.1

| *gSPL3.1 seq:* | TTAAAGGGGCATCAGTC\|CAATGGCAGAGAAATC | Mut | Genotype |
|---|---|---|---|
| Allele 1: | TTAAAGGGGCATCAGTC\|TCAATGGCAGAGAAATC | +1 | HM |
| Allele 2: | TTAAAGGGGCATCAGTC\|TCAATGGCAGAGAAATC | +1 | |

Nitab_0001538g0080.1

| *gSPL3.6 seq:* | CCTTCAACTTGACACCT\|AGGTGGCTGACCACCCT | Mut | Genotype |
|---|---|---|---|
| Allele 1: | CCTTCAACTTGACACCT\|AGGTGGCTGACCACCCT | WT | HT |
| Allele 2: | CCTTCAACTTG------\|--GTGGCTGACCACCCT | -8 | |

Nitab_0000991g0020.1

| *gSPL3.6 seq:* | CCTTCAACTTGACACCT\|AGGTGGCTGACCACCCTGT | Mut | Genotype |
|---|---|---|---|
| Allele 1: | CCTTCAACTTGACACCT\|AGGTGGCTGACCACCCTGT | WT | HT |
| Allele 2: | CCTTCAACTTGACACCT\|GTCTCGATTGATTGAATTCTG GTTCAAGTAAATTAGTGCTATCAGCGTACTTGGCATATG GCTATTATGAAACTTGGGTAGCACTTTGTCCATTTGAGT TAGTAGTTGGACTTTGGTATTGTATAATTAGGTGGCTGA CCACCCTGT | +123 | |

Nitab_0002299g0030.1

| *gSPL1.6 seq:* | ATGGACATAACAGGCGT\|CGAAGGAAACCTCAGCC | Mut | Genotype |
|---|---|---|---|
| Allele 1: | ATGGACATAACAGGCGT\|TCGAAGGAAACCTCAGCC | +1 | BA |
| Allele 2: | ATGGACATAACAGGCGT\|ACGAAGGAAACCTCAGCC | +1 | |

Nitab_0001010g0010.1

| *gSPL1.6 seq:* | ATGGACATAACAGGCGT\|CGAAGGAAACCTCAGCC | Mut | Genotype |
|---|---|---|---|
| Allele 1: | ATGGACATAACAGGCGT\|TCGAAGGAAACCTCAGCC | +1 | HM |
| Allele 2: | ATGGACATAACAGGCGT\|TCGAAGGAAACCTCAGCC | +1 | |

Nitab_0000067g0130.1

| *gSPL3.4 seq:* | CAGTCATTCCAAATGCC\|CAAAGGTCATCATAGGAGGTGCC | Mut | Genotype |
|---|---|---|---|
| Allele 1: | CAGTCATTCCAAAT--C\|CAAAGGTCATCATAGGAGGTGCC | -2 | HM |
| Allele 2: | CAGTCATTCCAAAT--C\|CAAAGGTCATCATAGGAGGTGCC | -2 | |

Nitab_0003900g0020.1

| *gSPL2.5 seq:* | GGAAACCATCCAGTTCA\|ACTCGGCAAGGCTTTCTTCAT | Mut | Genotype |
|---|---|---|---|
| Allele 1: | GGAAACCATCCAGTTCA\|ACTCGGCAAGGCTTTCTTCAT | WT | HT |
| Allele 2: | GGAAACCATCCAGTTCA\|AACTCGGCAAGGCTTTCTTCAT | +1 | |

Nitab_0000861g0050.1

| *gSPL2.4 seq:* | TGTGATCAGAGAGCCTC\|CTGCGGCAGCTTCTCTTCT | Mut | Genotype |
|---|---|---|---|
| Allele 1: | TGTGATCAGAGAGCCT-\|CTGCGGCAGCTTCTCTTCT | -1 | HM |

| | | |
|---|---|---|
| Allele 2: | TGTGATCAGAGAGCCT-\|CTGCGGCAGCTTCTCTTCT | -1 |

Nitab_0007217g0040.1

| *gSPL2.3 seq:* | TAGAAATTCACCCCCCA\|TGGAGGCTTTGGTAGGCT | Mut | Genotype |
|---|---|---|---|
| Allele 1: | TAGAAATTCACCCC---\|---------------GCT | -18 | HM |
| Allele 2: | TAGAAATTCACCCC---\|---------------GCT | -18 | |

**Plant SPL22-1**

Nitab_0003942g0050.1

| *gSPL3.3 seq:* | GAAATTCACAGACCTTG\|TGGCGGCGATGGTATGTC | Mut | Genotype |
|---|---|---|---|
| Allele 1: | GAAATTCACAGACCTTG\|TTGGCGGCGATGGTATGTC | +1 | BA |
| Allele 2: | GAAATTCACAGACCTT-\|TGGCGGCGATGGTATGTC | -1 | |
| *gSPL3.1 seq:* | TTAAAGGGGCATCAGTC\|CAATGGCAGAGAAATCAC | Mut | Genotype |
| Allele 1: | TTAAAGGGGCATCAGTC\|CAATGGCAGAGAAATCAC | WT | HT |
| Allele 2: | TTAAAGGGGCATCAGTC\|GCAATGGCAGAGAAATCAC | +1 | |

Nitab_0003348g0050.1

| *gSPL3.1 seq:* | TTAAAGGGGCATCAGTC\|CAATGGCAGAGAAATC | Mut | Genotype |
|---|---|---|---|
| Allele 1: | TTAAAGGGGCATCAGTC\|CAATGGCAGAGAAATC | WT | HT |
| Allele 2: | TTAAAGGGGCATCAGTC\|ACAATGGCAGAGAAATC | +1 | |

Nitab_0004959g0040.1

| *gSPL1.3 seq:* | AAGGGTCAAGTACTTCA\|ATGAGGTGTTGCCAAGCTGA | Mut | Genotype |
|---|---|---|---|
| Allele 1: | AAGGGTCAAGTACTTCA\|ATGAGGTGTTGCCAAGCTGA | WT | HT |
| Allele 2: | AAGGGTCAAGTACTTCA\|AATGAGGTGTTGCCAAGCTGA | +1 | |
| *gSPL1.4 seq:* | CGACGGCTTGAGCCTTT\|GCATGGTATTCACAGACTTTA | Mut | Genotype |
| Allele 1: | CGACGGCTTGAGCCTTT\|GCATGGTATTCACAGACTTTA | WT | HT |
| Allele 2: | CGACGGC----------------TATTCACAGACTTTA | -16 |  |

Nitab_0001538g0080.1

| *gSPL3.6 seq:* | CCTTCAACTTGACACCT\|AGGTGGCTGACCACCCT | Mut | Genotype |
|---|---|---|---|
| Allele 1: | CCTTCAACTTGACACCT\|AGGTGGCTGACCACCCT | WT | HT |
| Allele 2: | CCTTCAACTTGACACCT\|TAGGTGGCTGACCACCCT | +1 | |

Nitab_0002299g0030.1

| *gSPL1.6 seq:* | ATGGACATAACAGGCGT\|CGAAGGAAACCTCAGCC | Mut | Genotype |
|---|---|---|---|
| Allele 1: | ATGGACATAACAGGCGT\|ACGAAGGAAACCTCAGCC | +1 | BA |
| Allele 2: | ATGGACATAACAGGC-T\|CGAAGGAAACCTCAGCC | -1 | |

Nitab_0001010g0010.1

| *gSPL1.6 seq:* | ATGGACATAACAGGCGT\|CGAAGGAAACCTCAGCC | Mut | Genotype |
|---|---|---|---|
| Allele 1: | ATGGACATAACAGGCGT\|TCGAAGGAAACCTCAGCC | +1 | BA |
| Allele 2: | ATGGACATAACAGGCGT\|ACGAAGGAAACCTCAGCC | +1 | |

Nitab_0003900g0020.1

| *gSPL3.4 seq:* | CAGTCATTCCAAATGCC\|CAAAGGTCATTATAGCAGG | Mut | Genotype |
|---|---|---|---|
| Allele 1: | CAGTCATTCCAAATGCC\|CAAAGGTCATTATAGCAGG | WT | HT |
| Allele 2: | CAGTCATTCCAAATGC-\|CAAAGGTCATTATAGCAGG | -1 | |
| *gSPL2.5 seq:* | GGAAACCATCCAGTTCA\|ACTCGGCAAGGCTTTCTTCAT | Mut | Genotype |

| | | | |
|---|---|---|---|
| Allele 1: | GGAAACCATCCAGTTCA\|ACTCGGCAAGGCTTTCTTCAT | WT | HT |
| Allele 2: | GGAAACCATCCAGTTCA\|AACTCGGCAAGGCTTTCTTCAT | +1 | |
| *gSPL3.5 seq:* | AGATGGAGTGGAATGCA\|AAGTGGGACTGGGGAAACCTGG | Mut | Genotype |
| Allele 1: | AGATGGAGTGGAATGCA\|AAGTGGGACTGGGGAAACCTGG | WT | HT |
| Allele 2: | AGATGGAGTGGAAT---\|AAGTGGGACTGGGGAAACCTGG | -3 | |

Nitab_0000861g0050.1

| *gSPL2.4 seq:* | TGTGATCAGAGAGCCTC\|CTGCGGCAGCTTCTCTTCTTT | Mut | Genotype |
|---|---|---|---|
| Allele 1: | TGTGATCAGAGAGCCT-\|CTGCGGCAGCTTCTCTTCTTT | -1 | HM |
| Allele 2: | TGTGATCAGAGAGCCT-\|CTGCGGCAGCTTCTCTTCTTT | -1 | |
| *gSPL2.3 seq:* | TAGAAATTCACCCCCCA\|TGGAGGCTCCAGTAGGCTCTGT | Mut | Genotype |
| Allele 1: | TAGAAATTCACCCCCCA\|TGGAGGCTCCAGTAGGCTCTGT | WT | HT |
| Allele 2: | TAGAAATTCACCCCCCA\|ATGGAGGCTCCAGTAGGCTCTGT | +1 | |

**Plant SPL22-2**

Nitab_0003942g0050.1

| *gSPL3.3 seq:* | GAAATTCACAGACCTTG\|TGGCGGCGATGGTATGTC | Mut | Genotype |
|---|---|---|---|
| Allele 1: | GAAATTCACAGACCTTG\|TTGGCGGCGATGGTATGTC | +1 | BA |
| Allele 2: | GAAATTCACAGACCTT-\|TGGCGGCGATGGTATGTC | -1 | |
| *gSPL3.1 seq:* | TTAAAGGGGCATCAGTC\|CAATGGCAGAGAAATCAC | Mut | Genotype |
| Allele 1: | TTAAAGGGGCATCAGTC\|CAATGGCAGAGAAATCAC | WT | HT |
| Allele 2: | TTAAAGGGGCATCAGTC\|GCAATGGCAGAGAAATCAC | +1 | |

Nitab_0003348g0050.1

| *gSPL3.1 seq:* | TTAAAGGGGCATCAGTC\|CAATGGCAGAGAAATC | Mut | Genotype |
|---|---|---|---|
| Allele 1: | TTAAAGGGGCATCAGTC\|CAATGGCAGAGAAATC | WT | HT |
| Allele 2: | TTAAAGGGGCATCAGTC\|ACAATGGCAGAGAAATC | +1 | |

Nitab_0004959g0040.1

| *gSPL1.3 seq:* | AAGGGTCAAGTACTTCA\|ATGAGGTGTTGCCAAGCTGA | Mut | Genotype |
|---|---|---|---|
| Allele 1: | AAGGGTCAAGTACTTCA\|AATGAGGTGTTGCCAAGCTGA | +1 | HM |
| Allele 2: | AAGGGTCAAGTACTTCA\|AATGAGGTGTTGCCAAGCTGA | +1 | |
| *gSPL1.4 seq:* | CGACGGCTTGAGCCTTT\|GCATGGTATTCACAGACTTTA | Mut | Genotype |
| Allele 1: | CGACGGC----------------TATTCACAGACTTTA | -16 | HM |
| Allele 2: | CGACGGC----------------TATTCACAGACTTTA | -16 |  |

Nitab_0001538g0080.1

| *gSPL3.6 seq:* | CCTTCAACTTGACACCT\|AGGTGGCTGACCACCCT | Mut | Genotype |
|---|---|---|---|
| Allele 1: | CCTTCAACTTGACACCT\|AGGTGGCTGACCACCCT | WT | HT |
| Allele 2: | CCTTCAACTTGACACCT\|TAGGTGGCTGACCACCCT | +1 | |

Nitab_0002299g0030.1

| *gSPL1.6 seq:* | ATGGACATAACAGGCGT\|CGAAGGAAACCTCAGCC | Mut | Genotype |
|---|---|---|---|
| Allele 1: | ATGGACATAACAGGCGT\|ACGAAGGAAACCTCAGCC | +1 | BA |
| Allele 2: | ATGGACATAACAGGC-T\|CGAAGGAAACCTCAGCC | -1 | |

Nitab_0001010g0010.1

| *gSPL1.6 seq:* | ATGGACATAACAGGCGT\|CGAAGGAAACCTCAGCC | Mut | Genotype |
|---|---|---|---|
| Allele 1: | ATGGACATAACAGGCGT\|TCGAAGGAAACCTCAGCC | +1 | HM |
| Allele 2: | ATGGACATAACAGGCGT\|TCGAAGGAAACCTCAGCC | +1 | |

Nitab_0003900g0020.1

| *gSPL3.4 seq:* | CAGTCATTCCAAATGCC\|CAAAGGTCATTATAGCAGG | Mut | Genotype |
|---|---|---|---|
| Allele 1: | CAGTCATTCCAAATGCC\|CAAAGGTCATTATAGCAGG | WT | HT |
| Allele 2: | CAGTCATTCCAAATGC-\|CAAAGGTCATTATAGCAGG | -1 | |
| *gSPL2.5 seq:* | GGAAACCATCCAGTTCA\|ACTCGGCAAGGCTTTCTTCAT | Mut | Genotype |
| Allele 1: | GGAAACCATCCAGTTCA\|ACTCGGCAAGGCTTTCTTCAT | WT | HT |
| Allele 2: | GGAAACCATCCAGTTCA\|AACTCGGCAAGGCTTTCTTCAT | +1 | |
| *gSPL3.5 seq:* | AGATGGAGTGGAATGCA\|AAGTGGGACTGGGGAAACCTGG | Mut | Genotype |
| Allele 1: | AGATGGAGTGGAATGCA\|AAGTGGGACTGGGGAAACCTGG | WT | HT |
| Allele 2: | AGATGGAGTGGAAT---\|AAGTGGGACTGGGGAAACCTGG | -3 | |

Nitab_0000861g0050.1

| *gSPL2.4 seq:* | TGTGATCAGAGAGCCTC\|CTGCGGCAGCTTCTCTTCTTTT | Mut | Genotype |
|---|---|---|---|
| Allele 1: | TGTGATCAGAGAGCCT-\|CTGCGGCAGCTTCTCTTCTTTT | -1 | HM |
| Allele 2: | TGTGATCAGAGAGCCT-\|CTGCGGCAGCTTCTCTTCTTTT | -1 | |
| *gSPL2.3 seq:* | TAGAAATTCACCCCCCA\|TGGAGGCTCCAGTAGGCTCTGTT | Mut | Genotype |
| Allele 1: | TAGAAATTCACCCCCCA\|ATGGAGGCTCCAGTAGGCTCTGTT | +1 | HM |
| Allele 2: | TAGAAATTCACCCCCCA\|ATGGAGGCTCCAGTAGGCTCTGTT | +1 | |

**Plant SPL22-6**

Nitab_0003942g0050.1

| *gSPL3.3 seq:* | GAAATTCACAGACCTTG\|TGGCGGCGATGGTATGTC | Mut | Genotype |
|---|---|---|---|
| Allele 1: | GAAATTCACAGACCTT-\|TGGCGGCGATGGTATGTC | -1 | HM |
| Allele 2: | GAAATTCACAGACCTT-\|TGGCGGCGATGGTATGTC | -1 | |

Nitab_0003348g0050.1

| *gSPL3.1 seq:* | TTAAAGGGGCATCAGTC\|CAATGGCAGAGAAATC | Mut | Genotype |
|---|---|---|---|
| Allele 1: | TTAAAGGGGCATCAGTC\|ACAATGGCAGAGAAATC | +1 | HM |
| Allele 2: | TTAAAGGGGCATCAGTC\|ACAATGGCAGAGAAATC | +1 | |

Nitab_0004959g0040.1

| *gSPL1.3 seq:* | AAGGGTCAAGTACTTCA\|ATGAGGTGTTGCCAAGCTGA | Mut | Genotype |
|---|---|---|---|
| Allele 1: | AAGGGTCAAGTACTTCA\|AATGAGGTGTTGCCAAGCTGA | +1 | HM |
| Allele 2: | AAGGGTCAAGTACTTCA\|AATGAGGTGTTGCCAAGCTGA | +1 | |
| *gSPL1.4 seq:* | CGACGGCTTGAGCCTTT\|GCATGGTATTCACAGACTTTA | Mut | Genotype |
| Allele 1: | CGACGGC----------------TATTCACAGACTTTA | -16 | HM |
| Allele 2: | CGACGGC----------------TATTCACAGACTTTA | -16 |  |

Nitab_0001538g0080.1

| *gSPL3.6 seq:* | CCTTCAACTTGACACCT\|AGGTGGCTGACCACCCT | Mut | Genotype |
|---|---|---|---|
| Allele 1: | CCTTCAACTTGACACCT\|AGGTGGCTGACCACCCT | WT | HT |
| Allele 2: | CCTTCAACTTGACACCT\|TAGGTGGCTGACCACCCT | +1 | |

Nitab_0002299g0030.1

| *gSPL1.6 seq:* | ATGGACATAACAGGCGT\|CGAAGGAAACCTCAGCC | Mut | Genotype |
|---|---|---|---|
| Allele 1: | ATGGACATAACAGGCGT\|ACGAAGGAAACCTCAGCC | +1 | BA |
| Allele 2: | ATGGACATAACAGGC-T\|CGAAGGAAACCTCAGCC | -1 | |

Nitab_0001010g0010.1

| *gSPL1.6 seq:* | ATGGACATAACAGGCGT\|CGAAGGAAACCTCAGCC | Mut | Genotype |
|---|---|---|---|
| Allele 1: | ATGGACATAACAGGCGT\|ACGAAGGAAACCTCAGCC | +1 | HM |
| Allele 2: | ATGGACATAACAGGCGT\|ACGAAGGAAACCTCAGCC | +1 | |

Nitab_0003900g0020.1

| *gSPL3.4 seq:* | CAGTCATTCCAAATGCC\|CAAAGGTCATTATAGCAGG | Mut | Genotype |
|---|---|---|---|
| Allele 1: | CAGTCATTCCAAATGCC\|CAAAGGTCATTATAGCAGG | WT | HT |
| Allele 2: | CAGTCATTCCAAATGC-\|CAAAGGTCATTATAGCAGG | -1 | |
| *gSPL2.5 seq:* | GGAAACCATCCAGTTCA\|ACTCGGCAAGGCTTTCTTC | Mut | Genotype |
| Allele 1: | GGAAACCATCCAGTTCA\|ACTCGGCAAGGCTTTCTTC | WT | HT |
| Allele 2: | GGAAACCATCCAGTTCA\|AACTCGGCAAGGCTTTCTTC | +1 | |
| *gSPL3.5 seq:* | AGATGGAGTGGAATGCA\|AAGTGGGACTGGGGAAACC | Mut | Genotype |
| Allele 1: | AGATGGAGTGGAATGCA\|AAGTGGGACTGGGGAAACC | WT | HT |
| Allele 2: | AGATGGAGTGGAAT---\|AAGTGGGACTGGGGAAACC | -3 | |

Nitab_0000861g0050.1

| *gSPL2.4 seq:* | TGTGATCAGAGAGCCTC\|CTGCGGCAGCTTCTCTTCT | Mut | Genotype |
|---|---|---|---|
| Allele 1: | TGTGATCAGAGAGCCT-\|CTGCGGCAGCTTCTCTTCT | -1 | HM |
| Allele 2: | TGTGATCAGAGAGCCT-\|CTGCGGCAGCTTCTCTTCT | -1 | |
